# Agricultural drivers of field margin plant communities are scale dependent

**DOI:** 10.1101/2023.03.02.530797

**Authors:** Isis Poinas, Guillaume Fried, Laura Henckel, Christine N Meynard

## Abstract

In recent decades, agricultural intensification has led to a strong decline in biodiversity. Field margins act as shelters and dispersal corridors for biodiversity in highly disturbed landscapes, and are critical to the maintenance of ecosystem services. However, they are also impacted by agricultural practices in neighbouring fields. Agricultural impacts are often studied at field to landscape scales, and rarely across biogeographic regions. One of the challenges in large-scale studies is the lack of standardized monitoring schemes including both biodiversity and accurate estimation of agricultural practices. Here, we take advantage of a national monitoring scheme in 462 sites in France, to assess the effects of agricultural practices on field margin flora at different extents and resolutions. We used spatial simultaneous autoregressive and generalized dissimilarity models to assess the response of plant richness and composition to climatic, soil and landscape conditions, and to agricultural (fertilization, herbicides) and margin management drivers. Analyses were repeated at the site-level, 40 and 75 km resolutions, and at regional and national extents. We found that the impact of agricultural practices on species richness was most important at the site-level, whereas climate and crop diversity became more important at the 75 km resolution. Compositional variations responded differently, with climate being more important at the site-level, and fertilization and crop diversity at the coarsest resolution. There was a strong variation in the variance explained by models among regions, but climate effects were weaker within biogeographic units compared to the national level, and different agricultural practices stood out as influential in different regions, suggesting that the regional context is fundamental in determining plant community structure. To efficiently conserve biodiversity, we therefore recommend the implementation of agricultural measures adapted to each region.

## Introduction

Since the second half of the 20th century, agricultural intensification has resulted in steady declines in biodiversity across a wide range of taxa and habitats (Emmerson et al., 2016). The mechanization of agriculture and the resulting aggregation of cropped fields has contributed to the loss of semi-natural areas, such as herbaceous field margins, defined here as uncultivated vegetated strips located between the cropland and the adjacent habitat. Despite their modest area in agroecosystems, field margins play an important ecological role as sources of food, habitat and dispersal corridors for endangered species (Storkey et al., 2011), pollinators (Zamorano et al., 2020) and pest predators (Woodcock et al., 2016). Locally, they also buffer pesticide and fertilizer drift into adjacent terrestrial and aquatic habitats (Haddaway et al., 2018). Their importance has been increasingly recognized in recent European agricultural policies, for example through the conditioning of a green payment provided that margin strips are preserved (Matthews, 2013). Field margins also represent ecotones, with a mix of weed and ruderal species, which are well adapted to agricultural disturbances, along with some other species that are more typical of grasslands and forests (Aavik & Liira, 2009). All these characteristics make of field margins a valuable environment to study the unintended impacts of agriculture on biodiversity.

At the field scale, agricultural practices play a key role in shaping the species distribution and diversity of plants in field margins (Marshall & Moonen, 2002). Chemical inputs such as pesticides and fertilizers often drift to adjacent habitats creating unintended disturbances in field margin communities. Practices such as sowing date, fertilization level, herbicide use and management of the margin flora using mowing, herbicides or grazing, vary by crop and production type, selecting species that are adapted to these disturbances (Fried et al., 2018; Bassa et al., 2011). Agricultural practices also impact dispersal dynamics (Tscharntke et al., 2005). At the landscape scale, the proportion and diversity of non-crop habitats (Rader et al., 2014) and crop diversity (Sirami et al., 2019) have positive effects on field margin biodiversity. At the regional and national scale, cropping systems are defined by the sequence of crop successions along with the cultural practices associated to each crop (Leenhardt et al., 2010), which are themselves correlated to climate and topography for example. In France, this has been the basis to define stable uniform agricultural units called “agricultural regions” (Richard-Schott, 2009). Long-term practices over these homogeneous regions could potentially influence the available species pool at larger spatial and temporal scales. It is therefore possible that a given agricultural practice has different local effects on biodiversity depending on the regional species pool (Cornell & Harrison, 2014). Considering local and regional effects at different scales is therefore important to have a better understanding of agricultural practices that could favor biodiversity.

In ecology, spatial scale is usually defined by two attributes: resolution (or grain) and extent, both of them having a strong importance on the processes unveiled. Spatial grain is the size of the unit of analysis. It is often assumed that coarser resolutions allow ignoring local stochasticity to focus on the main patterns structuring communities (Chase, 2014). For example, landscape variables are expected to have the greatest influence when the grain size is closest to the average area of the different habitats. However, using excessively coarse resolutions may lead to uninformative patterns (Viana & Chase, 2019). Spatial extent, on the other hand, is the total area covered by the sampling. It determines the range of variability in some environmental gradients, as well as the potential of inclusion of dispersal processes (Viana & Chase, 2019). Naturally, local filters such as farming practices are easier to detect at small extents and within biogeographic regions, where different study sites are more likely to represent homogeneous conditions for large-scale gradients such as climate (Viana et al., 2016).

Few studies have examined the integration of several spatial scales in agroecosystems. Some have suggested that agricultural effects could prevail at fine spatial grains. For instance, Guerrero et al. (2014) found that weed species richness in Spain decreased linearly with field-level intensification, but remained constant at the landscape level. However, Berquer et al. (2021), focusing at a resolution under 1 km, found the opposite pattern, with field margin communities showing a greater influence of landscape than agricultural practices. Concepción et al. (2012) revealed that extensive farming has a weaker positive impact on weed diversity at the field center and inner field edge when the landscape is less diversified within a 500 m radius. This suggests that landscape-scale simplification or intensification may result in a depletion of the regional species pool, which then becomes primarily composed of species adapted to agricultural disturbances. Regarding spatial extent, although some studies have addressed national (Rader et al., 2014), continental (Billeter et al., 2008) or regional scales (Nascimbene et al., 2012), they have included a small sample size or have lacked standard survey protocols across plots, often using proxies rather than direct measures of the targeted agricultural practices. Therefore, issues of scale in agroecosystems remain largely under-represented in the literature.

Here we aimed at studying the effects of spatial resolution and extent, for a better understanding of the effects of agricultural practices on plant diversity and composition in field margins. We relied on a standardized national monitoring effort in 462 agricultural field margins covering France between 2013 and 2019, and including monitoring of plant communities as well as agricultural practices. Previous analyses of this dataset at a site-level showed that vegetation composition was primarily structured by landscape and soil, and secondarily by agricultural intensification (Andrade et al., 2021; Fried et al., 2018). Composition was driven by margin management and fertilization, while species richness responded to herbicide use. However, these previous analyses did not explore the role of scale in their results, or whether the relative influence of agricultural practices on margin plant communities were consistent across regions. Building on these first findings, here we investigated the potential influence of spatial extent and resolution of analysis on the drivers of field margin plant communities. Because agricultural practices are site-specific whereas landscape, soil and climatic conditions tend to have a greater range of values within larger grid cells, we hypothesized that the effects of the former would dominate at finer resolutions, while the latter would stand out at coarser resolutions. Regarding spatial extent, we expected soil and climate variables to dominate plant community structure at the national extent and become less important within biogeographic regions, providing a clearer picture of the influence of agricultural practices at the regional extent.

## Materials and methods

### Vegetation surveys

We used vegetation data from a national monitoring effort, the 500-ENI network, which is funded by the French Ministry of Agriculture (see details in Andrade et al., 2021). Note that raw data access requires a request to the Ministry and is conditional on confidentiality of some information, such as site coordinates; however, all datasets used in the following analyses are available in a repository (see the data accessibility statement). Agricultural field margins covering continental France, were surveyed yearly since 2013, representing a total of 543 unique sites between 2013 and 2019 (including some site turnover between years, see **Appendix A**, **Fig. SA.1**). Here we selected a subset of sites that had at least five years of botanical data between 2013 and 2019, resulting in a total of 462 sites for the rest of the analysis. These survey sites were located in field margins representing four main crop types: annual crops (with winter wheat or maize as the main crop production, but including other crops in the rotation), market gardening crops (mainly lettuce) and vineyards. The proportion of sites under organic farming was roughly 20%, but agricultural practices covered a wide range of pesticide application, fertilizers and soil management. Botanical surveys were performed at peak flowering (between the end of April and the beginning of August, depending on the region). At the national scale, this represented 3079 observations (year x site) and 689 plant species. The transect line within each site was located in the middle of the field margin, equidistant from the cropland and the adjacent habitat. Along this transect, plant species were identified in ten 1 m² sub-plots (**Appendix A**, **Fig. SA.2**). Presence-absence was recorded for each species and observation. Here we used frequency of occurrence averaged across years (0 = was never detected in that site; 1 = was registered for all surveyed years in that site), as an index of relative abundance in the species-by-site matrices.

### Explanatory variables

We gathered two sets of explanatory variables: the first set came directly from the 500-ENI network and reflects agricultural practices assessed directly on the sites (see Andrade et al., 2021); the second set was compiled from external open access databases. These include soil, climate and landscape data (see below).

Agricultural practices were reported yearly from interviews of farmers into an online database according to a national standardized acquisition protocol. This data related to fertilization, pesticide use, tillage and boundary management (**Appendix A**, **Fig. SA.3**). Soil characteristics in the 0-5 cm layer were acquired at a resolution of 250 m from SoilGrids (Hengl et al., 2014). Climatic data by year and season were obtained from the Chelsa database for the period 1979-2013, at a resolution of 1 km (Karger et al., 2017). By using this dataset, we assumed that climate did not change dramatically between this period and our sampling period, or that any change that did occur conserved a similar spatial structure of climatic variability across sites. We also extracted landscape descriptors from OSO land cover maps at a 10 m resolution for 2016 (Inglada et al., 2017). We considered percent cover of eight landscape classes (**Appendix A**, **Fig. SA.3**) within buffers of 1 km radius, and used the Shannon diversity index (SHDI) (Turner & Gardner, 2015) to characterize compositional landscape heterogeneity. A Shannon’s diversity index was also estimated for crop landscape diversity within a buffer of 1 km, considering 17 types of crops (**Appendix A**, **Fig. SA.3**) retrieved from the external databases OSO (Inglada et al., 2017), TOPO (IGN, 2019) and the Graphical Parcel Register (IGN & ASP, 2019).

Our dataset contained missing values for agricultural drivers that were occasionally left empty by the local observers. We used a multivariate imputation method based on a random forest algorithm (package mice of R v.4.0.0, Buuren & Groothuis-Oudshoorn, 2011) to complete these data (see **Appendix B** for details).

For compositional analyses, all explanatory variables were standardized to have equal mean and standard deviation. We limited the number of explanatory variables to seven (**Table 1**) so that we could include the same ones at different resolutions (see section on data analysis below). We aimed at representing the essential variables in different categories (climate, soil, landscape and practices) ensuring that they were not highly correlated (Pearson correlation coefficient < 0.6). We retained mean annual temperature and soil pH, since they are known to be tightly related to floristic composition and richness in agroecosystems (Fried et al., 2008). Landscape and crop diversity were selected as indicators of landscape heterogeneity (Rader et al., 2014; Sirami et al., 2019). Finally, we included the herbicide treatment frequency index, the nitrogen dose applied in fertilizers (within adjacent agricultural fields) and the number of margin management events (**Table 1**). These variables have consistently been reported to have a significant effect on field margin communities (Aavik & Liira, 2009; Bassa et al., 2011; Fried et al., 2018).

**Table 1.** Description of explanatory factors used in analyses, with their abbreviations and scale of computation. The variables computed by observation and then averaged by sites are underlined.

| Factors | Abbreviations | Units | Type of factor | Data source | Scale of computation |
| --- | --- | --- | --- | --- | --- |
| Mean annual temperature | TEMP | °C x 10 | Climate | Chelsa Climate | 1 km |
| Soil pH (water) | PH | pH x 10 | Soil | SoilGrids | 250 m |
| Shannon's habitat diversity index | SHDI |  | Landscape | OSO | 1 km |
| <u>Shannon's crop diversity index</u> | SHDI_C |  | Landscape | RPG | 1 km |
| <u>Dose of nitrogen (fertilization and amendments)</u> | N_DOSE | kg/ha | Agricultural practices | 500-ENI | Site |
| <u>Treatment Frequency Index of herbicides</u> | HERB |  | Agricultural practices | 500-ENI | Site |
| <u>Number of management events</u> | MAN |  | Margin management | 500-ENI | Site |
| Spatial structure | SPAT | m (Lambert 93) | Spatial | 500-ENI | Site |

### Data analysis

We used two statistical methods: spatial simultaneous autoregressive models (SAR) with species richness as response variable, and generalized dissimilarity models (GDM) to study species composition while dealing with nonlinear relationships between environmental distance and species turnover (Ferrier et al., 2007). These two types of analyses were applied at the national extent using different resolutions, and then at regional and national extents using the site-level resolution.

SAR models were implemented with the package spdep (function *errorsarlm*) using a correlated spatial error term, which is a function of distance between sites up to a threshold distance, which corresponds to the distance at which autocorrelation becomes non-significant (**Appendix C**) (Cressie, 2015). We can thus model linear relationships that take into account spatial autocorrelation in the data, i.e. the tendency of nearby points to have more similar values than expected by chance. Hereafter, the term “spatial structuring” will be used interchangeably with “spatial autocorrelation”, which can result from an effect of the environment or from an internal process linked to community assembly, such as dispersal limitations (Borcard et al., 2018). Species richness was transformed with a square root to comply with normality. We assessed the residual autocorrelation of each model with a Moran’s I test (Thioulouse et al., 2018). We also computed partial regressions using the Nagelkerke pseudo-R² (partial determination coefficient) to quantify the relative importance of each predictor (Lichstein et al., 2002; **Appendix C**).

The use of GDMs (R package gdm, function *gdm*) to explain variability in species composition is appropriate when species turnover responds non-linearly to environmental factors (Ferrier et al., 2007). The Bray-Curtis dissimilarity among all pairs of sites was modeled as a function of environmental distance (computed for each factor) and geographical distance, which accounts for spatial structure. The environmental table was permuted 500 times to assess variable significance, as recommended for analyses with distance matrices (Ferrier et al., 2007). We then evaluated the importance of each variable in the model with the same procedure as above (partial regressions based on the explained deviance; Maestri et al., 2017).

The different analyses were reiterated at different spatial scales. The first approach was to increase the study resolution by aggregating data into grid cells. We present here results from three resolutions: site-level, 40 km, and 75 km, (**Fig. 1A**, but see **Appendix D & E** for other resolutions). To mitigate the influence of outliers at aggregated scales, we excluded grid cells that contained all sites within less than 5 km from each other in the same grid cell. We also removed grid cells with only one site, as well as cells along land-sea borders with more than 50% of their surface in the water. We also shifted grid placement in three different iterations, to make sure results were robust to the grid placement (**Appendix D**). Overall, this led to some variations in the number of sites and grid cells across analyses, which were investigated by resampling (**Table 2**, **Appendix D**). Whenever possible, explanatory data were directly extracted at the scale of interest (climate, soil and landscape). All other factors were averaged by grid cell from site-level data, including the frequency of occurrence by site. Species richness in each grid cell was obtained by aggregating site-level species lists to recalculate richness at the aggregated scale. In all spatial analyses involving aggregated datasets, we used the coordinates of the grid cell centroids.

**Fig. 1.**
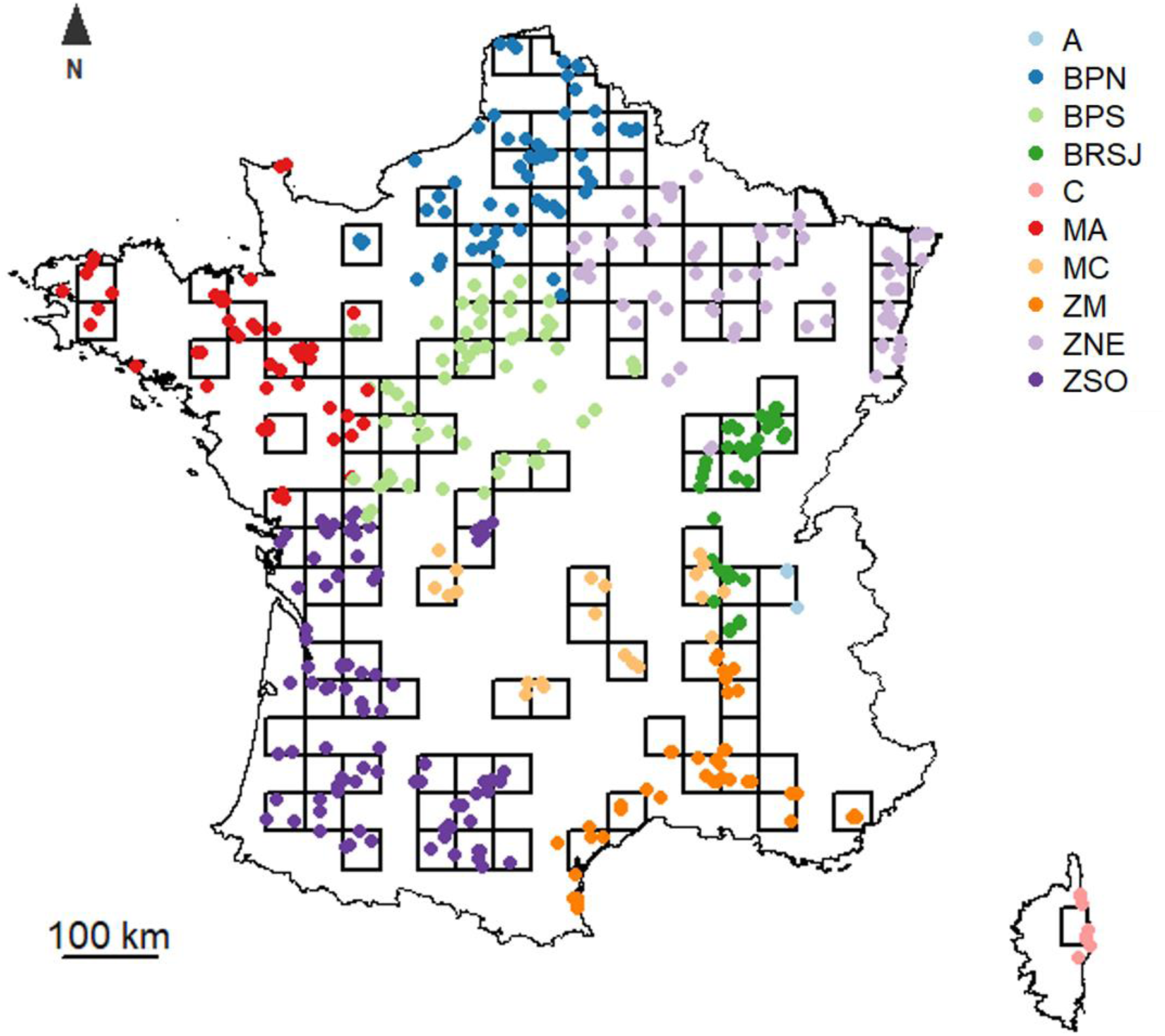
Distribution map of the field margins surveyed in France distributed in a grid of 40 km and colored according to their biogeographic region. Only cells selected for analyses are presented. A = « Alps », BPN = « Northern Parisian Basin », BPS = « Southern Parisian Basin », BRSJ = « Rhone-Saone-Jura Basin », C = « Corsica », MA = « Armorican Massif », MC = « Massif Central », ZM = « Mediterranean Zone », ZNE = « North-Eastern France » and ZSO = « South-Western France ». Note that the regions C, A, MC and BRSJ have been removed from analyses because of their small number of sites.

**Table 2.**
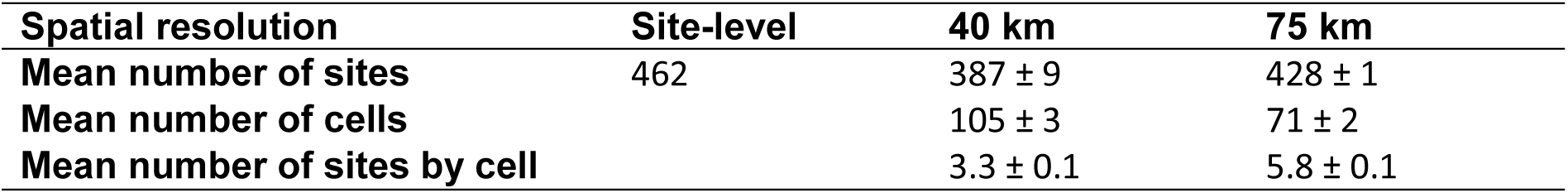
Sample size for each resolution. Grid positions in the aggregated resolutions were shifted to obtain three different positions of the grid system. Means and standard deviations below reflect variations across these grids within each resolution.

| **Spatial resolution** | **Site-level** | **40 km** | **75 km** |
| --- | --- | --- | --- |
| **Mean number of sites** | 462 | 387 ± 9 | 428 ± 1 |
| **Mean number of cells** |  | 105 ± 3 | 71 ± 2 |
| **Mean number of sites by cell** |  | 3.3 ± 0.1 | 5.8 ± 0.1 |

Lastly, we carried out the same analysis using site-level data at two different extents: national and regional. For this purpose, sites were grouped within ten biogeographical regions standing for the wide pedo-climatic and floristic gradients in France (**Fig. 1B**, based on the VégétalLocal map; Office français de la biodiversité, 2021). Only six regions had a sufficient number of sampling sites to be considered in our analyses (>40 sites): Northern Parisian Basin, Southern Parisian Basin, Armorican Massif, Mediterranean Zone, North-Eastern France and South-Western France.

## Results

### Effects of spatial resolution

At the site-level resolution, the predictors explained 27.8% of the variance in species richness. Herbicides and nitrogen dose were the main drivers and explained respectively 1.5 (p = 0.002) and 1.3% (p = 0.004) of the total variance in species richness, both having a negative effect on richness (**Table 3A**). There was a positive correlation of margin management with species richness that was mainly apparent at the site-resolution (p = 0.018). With respect to plant community composition, the GDM explained 29.5% of deviance (**Table 3B**), the main drivers being mean annual temperature (6.9%, p < 0.001) and soil pH (1.2%, p < 0.001).

**Table 3.** Significance of each predictor (in row) for each spatial resolution (in column, in km). For species richness in A), the direction of the relationship is indicated by +/-. The number of +/-/* indicates the number of times the predictor was significant over the three grid shifts. We only report significant relationships with p-values in brackets.

**A) Species richness**
|  | **0** | **40** | **75** |
| --- | --- | --- | --- |
| **Temperature** | + (0.004) | + (0.010) | ++ (< 0.001;<br>0.022) |
| **Ph** |  |  |  |
| **Landscape diversity** |  |  |  |
| **Crop diversity** |  |  | +++ (0.013;<br>0.007;<br>0.030) |
| **Herbicides** | - (0.002) | - (0.031) |  |
| **Nitrogen dose** | - (0.004) |  | - (0.044) |
| **Margin management** | + (0.018) | + (0.019) |  |
| **Spatial structure** | (< 0.001) |  |  |

**B) Composition**
|  | **0** | **40** | **75** |
| --- | --- | --- | --- |
| **Temperature** | (< 0.001) | *** (< 0.001;<br>< 0.001;<br>< 0.001) | *** (0.020;<br>< 0.001;<br>0.040) |
| **Ph** | (< 0.001) |  |  |
| **Landscape diversity** | (< 0.001) |  |  |
| **Crop diversity** | (< 0.001) | ** (< 0.001;<br>< 0.001) | *** (< 0.001;<br>0.040;<br>0.020) |
| **Herbicides** |  |  |  |
| **Nitrogen dose** |  | ** (< 0.001;<br>< 0.001) | ** (0.020;<br>< 0.001) |
| **Margin management** |  | * (< 0.001) |  |
| **Spatial structure** | (< 0.001) | *** (< 0.001;<br>< 0.001;<br>< 0.001) | *** (< 0.001;<br>< 0.001) |

With increasing spatial resolution, the importance of climate and landscape (i.e. crop diversity) for species richness increased, while agricultural effects remained constant and small (**Fig. 2A-C**). Temperature had a positive correlation with richness (p = 0.004), while soil pH had no significant effect. Crop diversity was the predominant landscape factor at all aggregated scales, showing a consistent positive correlation with richness, but only significant at the coarsest resolution (75 km, p = 0.017, **Table 3A**). On the contrary, landscape diversity did not seem to have any effect (**Table 3A**). Both herbicides and nitrogen dose reduced the number of species at the site-level, but these effects were not significant at coarser resolutions (**Table 3A**). The proportion of richness that can be explained by spatial autocorrelation was higher at the site-resolution (11.4%, p < 0.001) compared to coarser resolutions where it was non-significant (**Fig. 2A-C**).

**Fig. 2.**
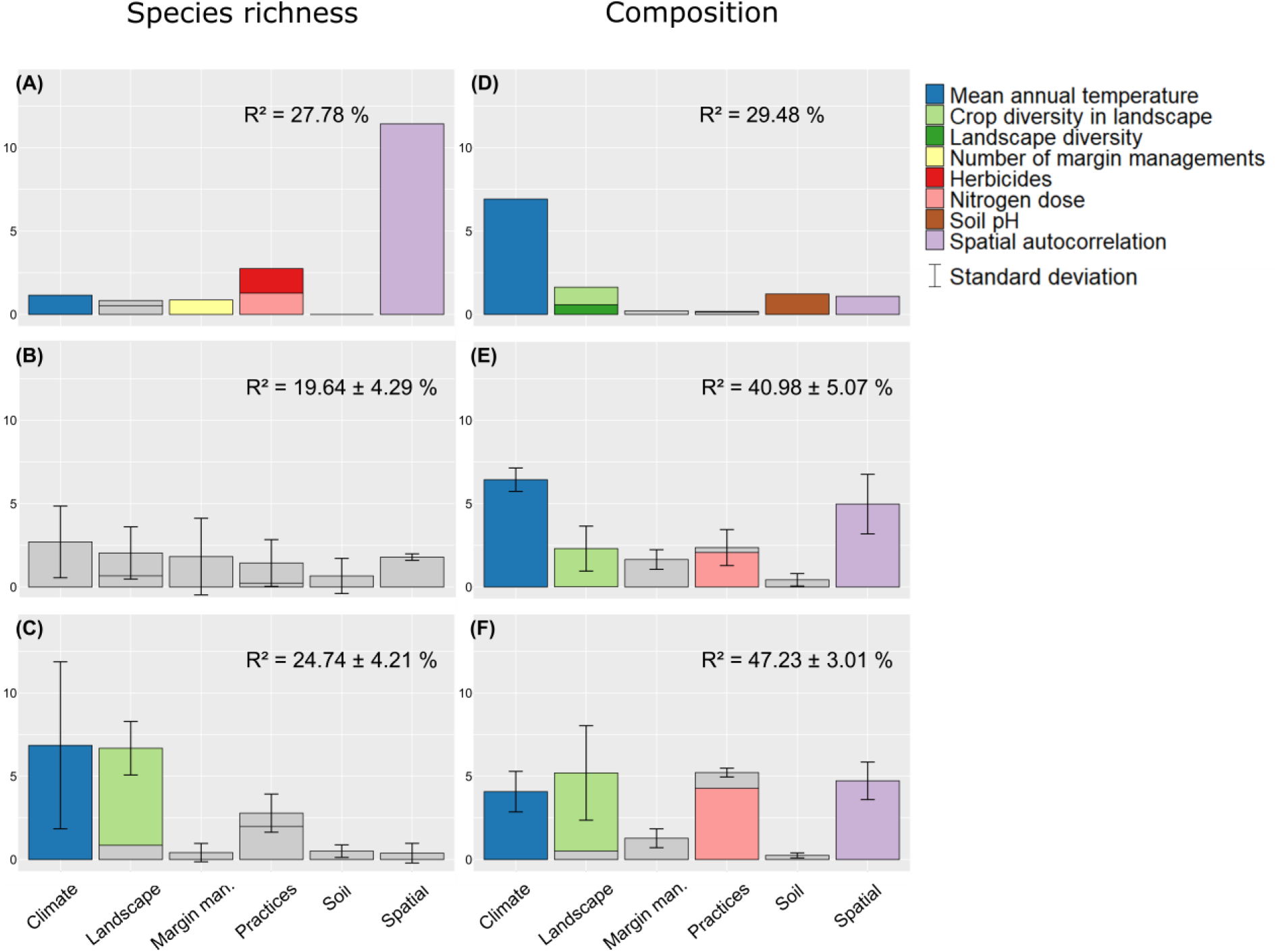
Percentage of explained variance of species richness and composition by each factor for each spatial resolution. Factors significant in less that one grid are in grey. (A, D) 0 km; (B, E) 40 km; (C, F) 75 km. Standard deviations are computed from the three resamplings of grids. Percentages of explained variance are reported in **Appendix E, Table SE.1**.

With regard to composition, the R² of the models increased at coarser spatial grains (**Appendix E**, **Fig. SE.1**). Climate and landscape (mainly through crop diversity) together contributed to overall differences in species composition among sites, much more strongly than farming practices and soil at all scales (**Fig. 2D-F**). Climate had less effect on composition at coarser resolutions while the effect of landscape increased. Fertilization was more strongly related to species composition at the coarsest resolution (75 km, 4.3%, p = 0.010) than at the finest, for which it was not significant (**Fig. 2D-F**, **Table 3B**). Herbicides did not have any effect on species composition. Also, spatial autocorrelation was relatively important at all aggregated scales (**Fig. 2E-F**).

### Effects of spatial extent

We compared the outcomes between national and regional extents across different regions. At the regional extents, the influence of environmental filters on species richness and composition varied greatly among regions (**Fig. 3**). For example, the high R^2^ of the model for the Mediterranean zone (approximately 38.8%) contrasted with the lower one for the Armorican Massif (15.1%).

**Fig. 3.**
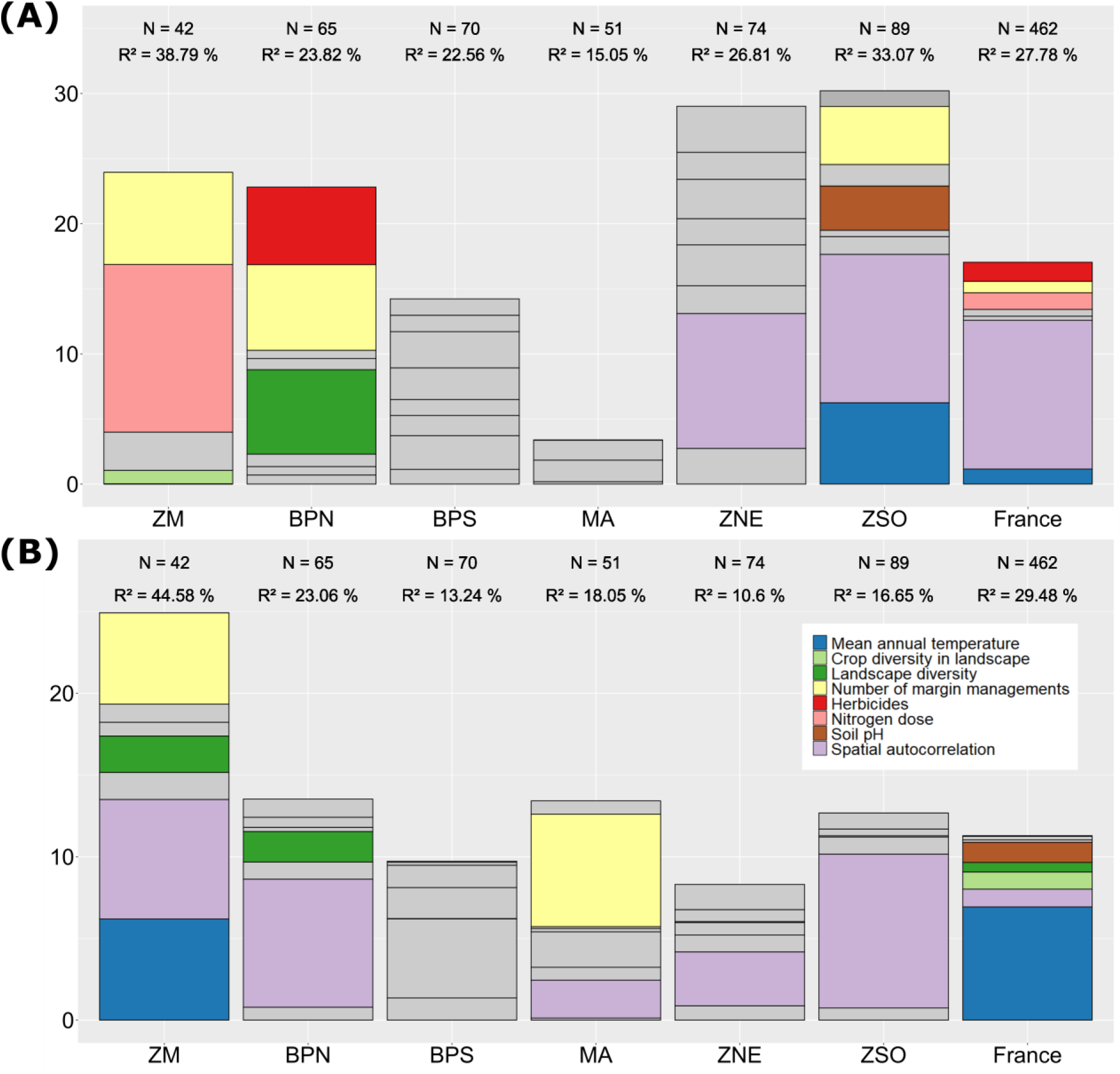
Percentage of explained variance at the site-scale resolution for (A) richness and (B) composition for each biogeographic region and national extent. Non-significant factors are in grey. Regions are ordered by area from left to right. Percentages of explained variance are reported in **Appendix E, Table SE.2**.

Some landscape and agricultural effects emerged at a regional extent as more significant than at the national extent. For example, landscape diversity was negatively correlated with richness (6.5%, p = 0.016) in the northern Parisian Basin, a relationship that was not visible at the national extent with all regions analyzed together (**Table 4A**). Agricultural effects on richness were often more predominant at the regional extent (**Fig. 3A**). This scale allowed for a clearer identification of the effects of nitrogen fertilization, herbicides, and margin management. The direction of the relationships between richness and predictors remained consistent with analyses at the national level, except for south-western France, which depicted a negative relationship of species richness with temperature, (p = 0.003, **Table 4A**).

**Table 4.** Significance of each predictor (in row) for each biogeographic region and for national extent (in column, see **Fig. 1** for abbreviations of region names). For species richness in A), the direction of the relationship is indicated by +/-. We only report significant relationships with p-values in brackets.

**A) Species richness**
|  | BPN | BPS | MA | ZM | ZNE | ZSO | France |
| --- | --- | --- | --- | --- | --- | --- | --- |
| Temperature |  |  |  |  |  | - (0.003) | + (0.004) |
| Ph |  |  |  |  |  | - (0.048) |  |
| Landscape diversity | - (0.016) |  |  |  |  |  |  |
| Crop diversity |  |  |  | + (0.021) |  |  |  |
| Herbicides | - (0.034) |  |  |  |  |  | - (0.002) |
| Nitrogen dose |  |  |  | - (0.001) |  |  | - (0.004) |
| Margin management | + (0.023) |  |  | + (0.017) |  | + (0.013) | + (0.018) |
| Spatial structure |  |  |  |  | (0.003) | (0.002) | (< 0.001) |

**B) Composition**
|  | BPN | BPS | MA | ZM | ZNE | ZSO | France |
| --- | --- | --- | --- | --- | --- | --- | --- |
| Temperature |  |  |  | (< 0.001) |  |  | (< 0.001) |
| Ph |  |  |  |  |  |  | (< 0.001) |
| Landscape diversity | 0.040 |  |  | 0.020 |  |  | (< 0.001) |
| Crop diversity |  |  |  |  |  |  | (< 0.001) |
| Herbicides |  |  |  |  |  |  |  |
| Nitrogen dose |  |  |  |  |  |  |  |
| Margin management |  |  | (< 0.001) | (< 0.001) |  |  |  |
| Spatial structure | (< 0.001) |  | (< 0.001) | (< 0.001) | (0.040) | (< 0.001) | (< 0.001) |

With respect to composition, climatic conditions appeared to have higher explanatory values at a national rather than regional extent (**Fig. 3B**). As with richness, new and contrasting effects of landscape and practices were highlighted within regions, such as the effect of landscape diversity on composition in the Mediterranean zone (2.2%, p = 0.020), and the effects of the number of margin management events in the Mediterranean zone (5.6%, p < 0.001) and Armorican Massif (6.9%, p < 0.001) (**Fig. 3B**, **Table 4B**).

## Discussion

Our results reveal the importance of scale, both resolution and extent, in understanding the non-intended effects of agricultural practices on field margin vegetation. In line with our expectations, the effects of agricultural practices on field margin flora were more prominent within biogeographic regions than when analyzed at the national extent. Herbicides and fertilization were important to explain species richness at the site-level, whereas fertilization stood out in explaining species composition at coarse resolutions. Conversely, climate and landscape variables always dominated when data were aggregated, and when analyzed at the national extent. Climate also explained a large percentage of variation in species composition, but not in species richness, at the site-level.

### Influence of spatial resolution

Interestingly, richness and composition did not respond to the same environmental variables. Richness was mostly impacted by local agricultural intensification, such as high herbicide use and nitrogen fertilization, while composition was influenced by large-scale landscape and agricultural effects, such as landscape crop diversity and high regional fertilization. This is consistent with some studies reporting that the effects of local agricultural practices on field margin plant communities are at least as important as landscape effects (Bassa et al., 2011; Gabriel et al., 2006; but see Martin et al., 2020). Negative impacts of herbicides and nitrogen on field margin richness have already been reported (Aavik & Liira, 2009; Fried et al., 2018), with nitrogen fertilization affecting both richness and composition, and herbicides only impacting richness (Fried et al., 2018). Our results revealed a large-scale effect of nitrogen use on composition, which contrasts with previous studies showing its effect on richness (Billeter et al., 2008; Kleijn et al., 2009). Storkey et al. (2011) demonstrated that large-scale eutrophication is a major explanatory factor of the threat status of arable species in Europe and even more than herbicide use, although the two factors are difficult to disentangle.

Regarding landscape variables, crop diversity was the most important predictor of richness and composition at large scales. While the response of plant species richness to Shannon crop diversity is inconsistent among studies (Martin et al., 2020; Sirami et al., 2019), we found here a consistently positive effect regardless of the scale. This can be viewed as landscape complementation for species that are specialists of a single crop (Storkey et al., 2011). Another explanation would be that crop diversity is negatively related to field size (Martin et al., 2020), and this latter might have an additional effect on richness (Fahrig et al., 2015). Furthermore, crop diversity may reflect the varied environmental conditions within a grid cell, and it is possible that species richness exhibits a stronger response to this environmental diversity. In contrast, habitat diversity did not show any effect on richness, but only on composition at the site level. Although a diverse landscape does not necessarily support more species, it may favor functionally more diverse communities, supporting the idea that species richness alone cannot capture all aspects of biodiversity (Aavik et al., 2008; Fried et al., 2018). The effect of crop diversity on composition increased with resolution, while this was not the case for landscape diversity, likely because landscapes became more homogeneous in terms of habitat diversity, while they still remained different in terms of crops (**Appendix F**, **Fig. SF.1**).

### Influence of spatial extent

More interestingly, our results pointed to strong regional specificities, underlining complex interactions between local and regional scales. For example, the influence of landscape diversity on richness was very high in the northern Parisian Basin, a region mainly composed of large cereal open-fields (**Appendix F**, **Fig. SF.2-3**). The direction of this effect was unexpected, since landscape diversity decreased the number of species in field margins. Given the high level of local intensification in this region, the species from natural habitats are likely too vulnerable to establish locally. Because diverse landscapes have fewer croplands and more natural habitats, species adapted to agricultural disturbances could be less prevalent in the landscape’ species pool, ultimately leading to a loss of species in highly disturbed sites. This apparently counter-intuitive finding could thus be explained by interactions between the regional pool and local conditions (Cornell & Harrison, 2014; Myers & Harms, 2009). Likewise, the detrimental effect of nitrogen fertilization on richness was only significant in the Mediterranean Zone. In this region, soils had low nitrogen content (**Appendix F**, **Fig. SF.3**), and the species pool is mostly composed of stress-tolerant species adapted to nutrient-poor soil conditions. Due to its regional composition, Mediterranean flora would thus be more sensitive to soil enrichment. This supports the results of Kleijn et al. (2009) who concluded that the impact of nitrogen is greater in less intensive regions. Clarifying regional specificities should thus be a strong priority for future agroecological studies, as it paves the way for proposing concrete agricultural management practices that minimize impacts on biodiversity while considering regional characteristics.

### Implications for management in agricultural context

Our results have practical implications for crop management. Crop diversity stands out with a major positive impact on plant diversity in field margins. The importance of crop diversity has just begun to be recognized in public policies (Galán-Martín et al., 2015), and our results suggest that this effect will extend to a large scale. Promoting a wide range of crops at the landscape level could therefore slow the decline of some highly specialized arable plant species, particularly in landscapes with large semi-natural cover (Sirami et al., 2019). Fertilization significantly altered floristic assemblages and is also more impactful at a large scale, which is of major interest for national agricultural policies.

Beyond large-scale agricultural impacts, the effects of local agricultural practices are also regionally dependent. The region in which regulatory or management measures are implemented can determine their cost-benefit (Kleijn et al., 2009). For instance, a drastic reduction of fertilizers in highly intensive regions mainly composed of nitrophilous species, such as the Parisian Basin here, will have little impact on plant communities of field margins if this action is not coupled with landscape restoration. In contrast, implementing stricter fertilization control in the Mediterranean zone, where plants are adapted to poor soils, would have a much higher impact on the native flora of this region. Regions that still have a diverse pool of species (such as the Mediterranean one) are thus more susceptible to respond to less costly mitigation measures and are of higher conservation concern (Stevens et al., 2010). We therefore encourage agricultural stakeholders and scientists involved in monitoring programs to further evaluate the changes in field margin flora and its protection measures across different spatial scales, and implement regulatory measures that incorporate landscape-level management planning adapted to each biogeographic and agricultural region.

## Acknowledgements

The 500-ENI network is supported by the French Ministry of Agriculture under the Ecophyto framework with funding from the French Biodiversity Agency (Office Français de la Biodiversité). We would like to thank everyone that has collected data in the field, the farmers who provided information on their practices, and everyone involved in the coordination of the 500-ENI data network.

## Authorship statement

I.P., G.F. and C.M. planned and designed the research; L.H. contributed to data processing; I.P. analyzed the data and wrote the first draft of the manuscript; G.F., L.H. and C.M. contributed substantially to revisions. All authors gave final approval for publication.

## Data accessibility statement

Data available via the Data INRAE Repository, at https://doi.org/10.57745/NMIPCY

## Appendix A-F. Supplementary data

Supplementary data associated with this article can be found, in the online version, at XXXXX.

