## Appendix for "Agricultural drivers of field margin plant communities are scale dependent"

**Appendix A.** Details about the sampling protocol and available data in the 500-ENI network.

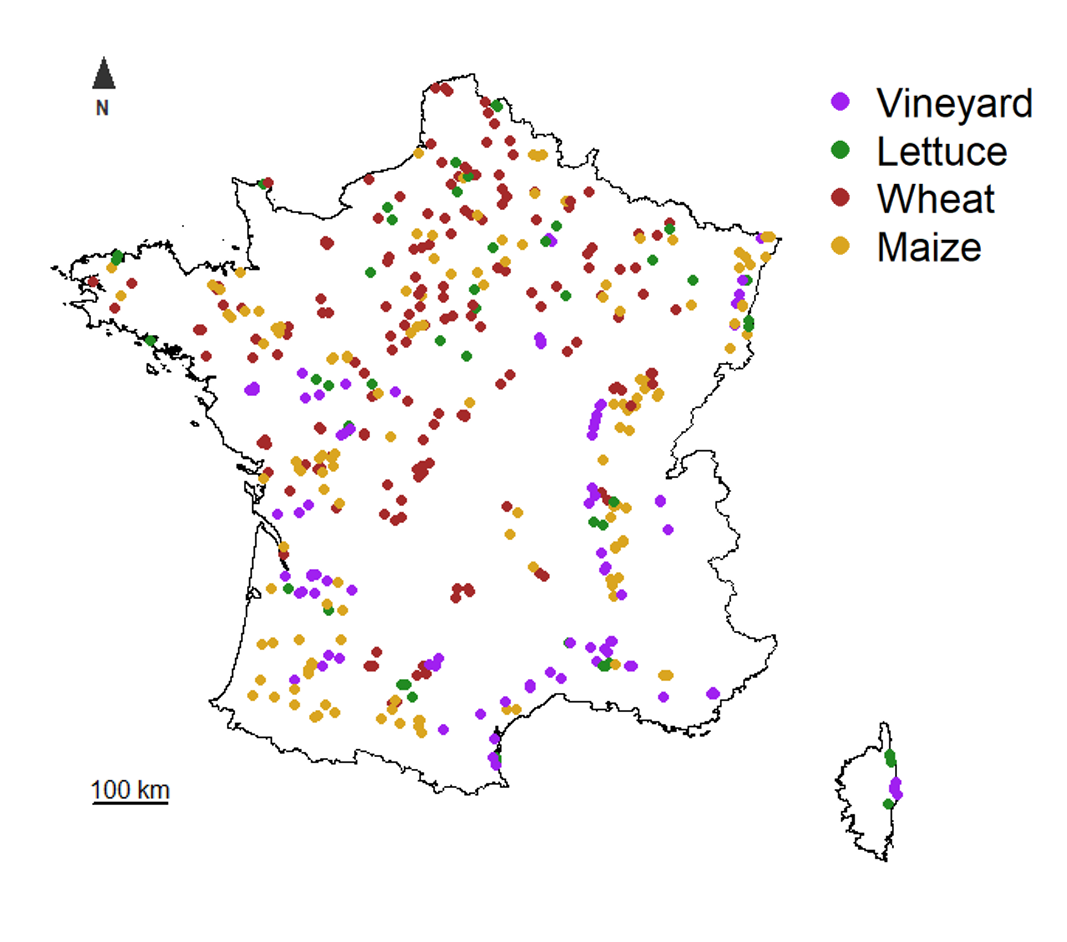

**Fig. A.1.** Distribution of fields monitored at least five years between 2013 and 2019 in continental France. Purple: vineyards (n = 93), green: market gardening crops (n = 50), brown: winter wheat in rotation (n = 178), yellow: maize in rotation (n = 141).

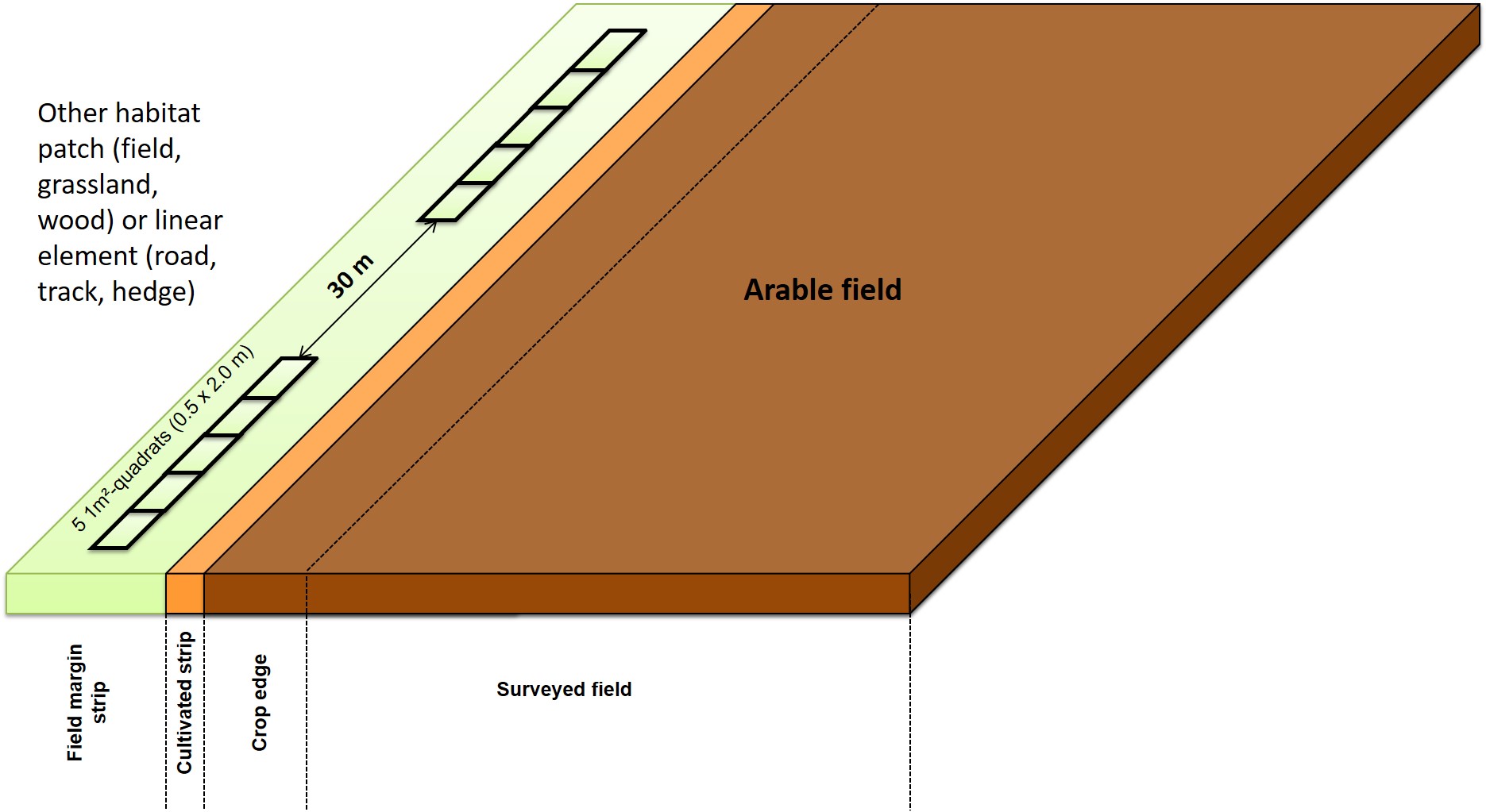

**Fig. A.2.** Details about the sampling protocol within each field margin (Andrade et al., 2021). The *crop edge* is the 1-6 first meters of the crop. The *cultivated strip* or *crop strip* is outside the last row of crops and is mostly composed of bare soil usually colonized by weed species from the field. The *field margin strip* is the uncultivated herbaceous strip between the *cultivated strip* and the *adjacent habitat*.

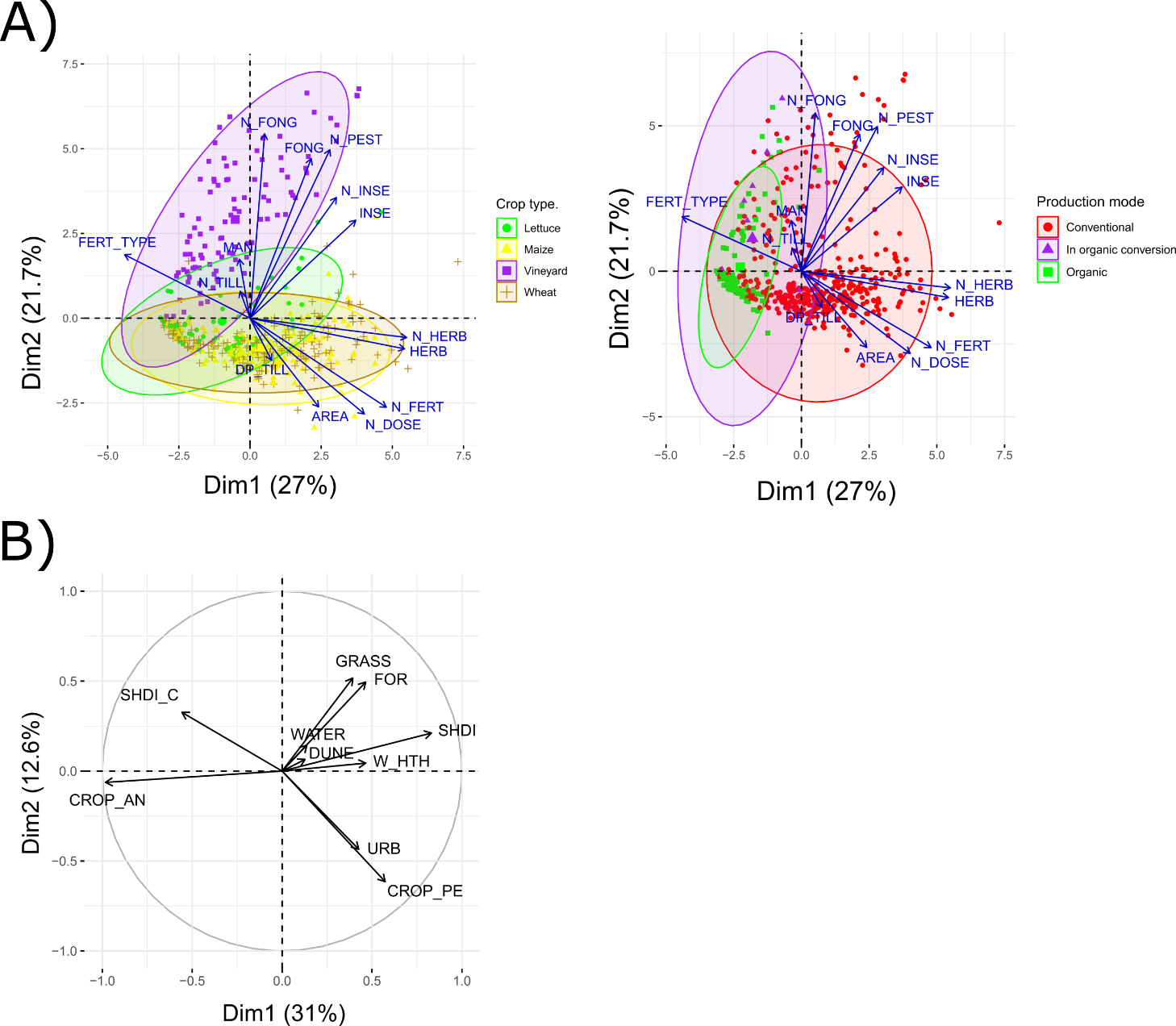

**Fig. A.3.** PCA on the main types of factors. Points stand for the 543 sites. A) Agricultural practices, B) Landscape. In A), agricultural practices are discriminated by crop type and production mode depicted by colored ellipses. For landscape, the SHDI was computed on 8 habitat classes (annual crops, perennial crops, forests, urban, grasslands, water, beaches and dunes, woody heaths) and the SHDI_C on 17 crop groups (maize, winter cereals, spring cereals, winter rapeseed, spring rapeseed, sunflower, beet, oilseed, grapevine, flax, arboriculture, fiber plants, vegetables or flowers, aromatic/medicinal/perfume plants, other crops, fallows and grasslands).

**Abbreviations for agricultural practices**

**AREA =** Field area

**DP_TILL =** Maximum depth of tillage

**FERT_TYPE =** Type of fertilization (mineral VS organic)

**FONG =** Fungicide treatment frequency index

**HERB =** Herbicide treatment frequency index

**INSE =** Insecticide treatment frequency index

**MAN =** Number of margin managements

**N_DOSE =** Nitrogen dose (fertilization and amendments)

**N_FERT =** Number of fertilizations and amendments

**N_FONG =** Number of fungicide applications

**N_HERB =** Number of herbicide applications

**N_INSE =** Number of insecticide applications

**N_PEST =** Number of pesticide applications

**N_TILL =** Number of tillage events

**Abbreviations for landscape**

**CROP_AN =** Annual crops

**CROP_PE =** Perennial crops (vineyards and orchards)

**DUNE =** Beaches and dunes

**FOR =** Forests

**GRASS =** Grasslands and lawns

**SHDI =** Shannon landscape diversity index

**SHDI_C =** Shannon crops diversity index

**URB =** Urban, roads, mineral surfaces

**W_HTH =** Woody heaths

**WATER =** Streams, rivers, lakes

**Table A.1.** Details about calculation of each explanatory factor included in analyses (see **Table 1**).

| Factors | Calculation |
| --- | --- |
| Mean annual temperature |  |
| Soil pH (water) |  |
| Shannon’s habitat diversity index | $- \sum_{i=1}^{n} p_{i}\ln p_{i}$  With $p_{i}$the proportion of the habitat i in the landscape |
| Shannon’s crop diversity index | $- \sum_{i=1}^{n} p_{i}\ln p_{i}$  With $p_{i}$the proportion of crop i in the agricultural area |
| Dose of nitrogen (fertilization and amendments) | $\frac{Quantity\times N of the formulation}{100}$ |
| Treatment Frequency Index of herbicides | $\frac{Applied dose of used products\times treated area}{certified dose\times plot area}$  For herbicides in vineyards, the treated area accounts for one third of the plot area if the treatment is done on the rows and two thirds if it is done on the inter-rows. In case of missing information, the treatment is supposed to involves only the rows (i.e. the most frequent practice). |
| Number of management events | All types of management (mowing, grazing…) |
| Spatial structure | Based on geographical coordinates, see **Appendix C** for more details |

**Appendix B.**  Method of imputation for missing values.

We used the R package mice (function of the same name) to impute the missing values from our dataset. This method relates on multiple imputations by chained equations in which each incomplete variable is imputed by a separate model. The algorithm imputes an incomplete variable using other variables as predictors. Variables are imputed in a random order. If predictors are incomplete themselves, the most recently generated imputations are used to complete the predictors prior to imputation of the target variable. The number of iterations is the number of times each variable is computed (until convergence). Given the large number of incomplete predictors in our dataset, we fixed this number to 10 (instead of 5 by default). The function offers a wide range of available methods depending on the nature of the data to be imputed. Here, we chose the Random forest algorithm, as a reliable method to avoid over-fitting by ensemble learning (Shah et al., 2014). We imputed the missing data 100 times to control for the variability of imputations. The ultimate imputed value for each observation is the one that occurs most frequently among the 100 imputations. To further improve the imputation, we included some complete factors not used in the analyses (climate, soil, landscape, adjacent habitat, width margin, field area, geographical coordinates, and all practices including the total number of fertilizations, insecticide, herbicide and fungicide applications, tillage and management events one year before the observation). The final dataset given in input included 44 variables and 13119 observations (observations from other groups than plants were added to improve the learning ability of the algorithm).

The robustness of the method has been tested for 3013 observations. We ensured that the percentage of missing values in each factor did not change from the full dataset. We checked the differences between observed and imputed values. We saw that the error rate of a variable is partly correlated to its percentage of missing values but never exceeds 10% of the total values imputed, and 2.2% of the total dataset (3013 observations).

We also ensured that the imputations by observation did not change significantly the mean values by site. We focused on a subset of 58 sites for which we were able to compute the difference in percentage of the imputed value with the correct value (sites for which at least 5 years of data were available for all variables of interest). It results in 91% of the sites having an estimate of nitrogen dose within 10% of the correct value (100% of the sites for other variables). When differences were tested over the whole dataset (means before VS after imputation), it results in a change of less than 25% from the original value for 98% of the sites, except for nitrogen dose for which it was 76% of sites.

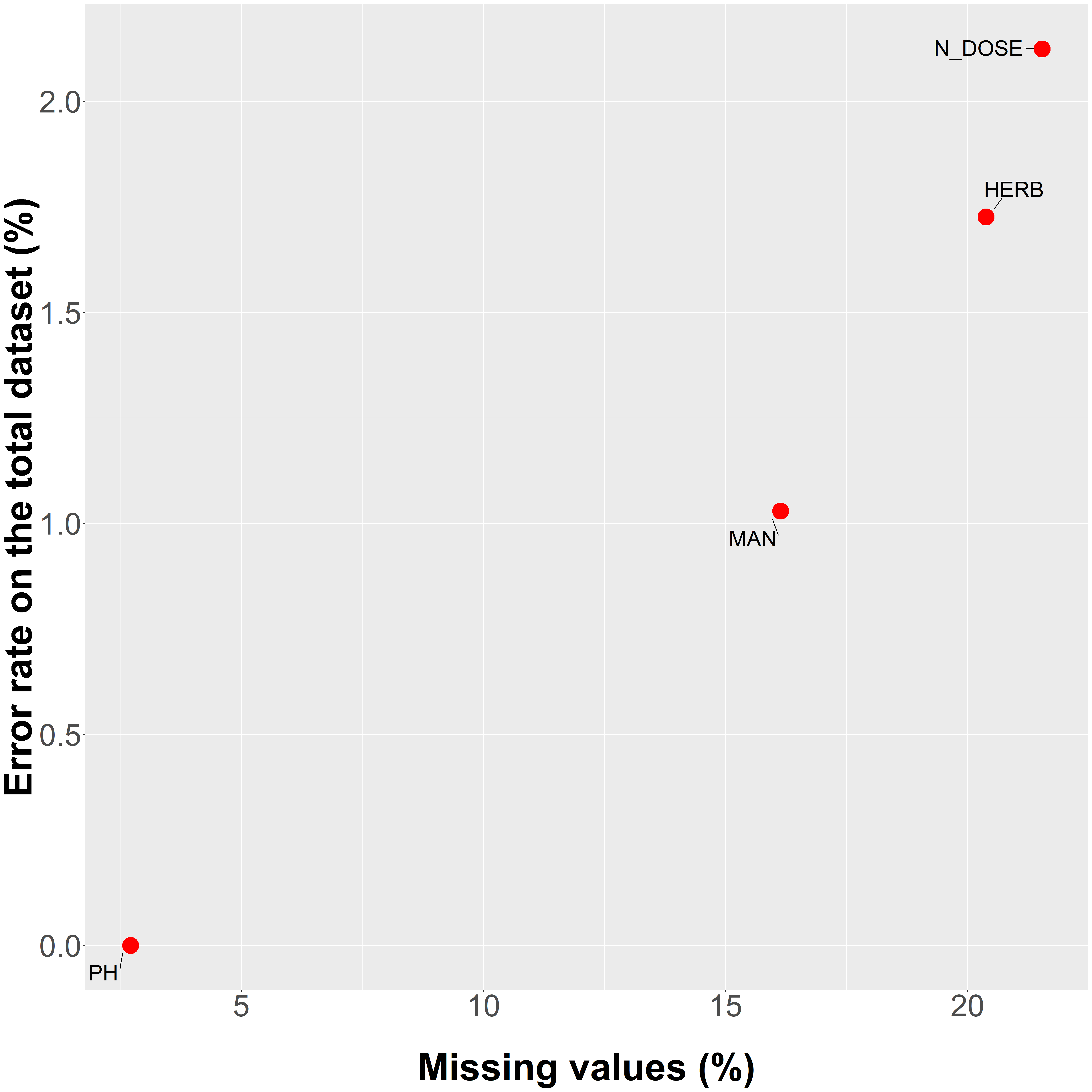

**Fig. B.1.** Error rate of imputed values estimated for the total used dataset according to the percentage of missing values of each variable.

**Appendix C.** Spatial parameters for SAR and methodology for partial regression analyses.

Spatial correlograms were used to choose the maximal distance for the spatial neighborhood matrix at each resolution. We tested several distances around the visual estimate for each spatial resolution and selected the one that produced the lower AIC in a spatial regression with no other predictors (Kissling & Carl, 2008; **Table SC.1**). Similarly, we used AIC to select one of two weight functions (1/x and 1/x²) for the neighborhood matrix.

**Table C.1.** Maximum distances for spatial neighborhood matrix. The selected weight function was always 1/x.

|  | 0 | 25 | 40 | 60 | 75 |
| --- | --- | --- | --- | --- | --- |
| Maximum distance (in km) | 25 | 25 | 40 | 80 | 100 |

We followed the methodology described in Borcard *et al.* (1992) to partition the R² of the model in several parts, each of them attributed to a single explanatory variable or to the spatial component itself (spatial autocorrelation). For GDM, we used the explained deviance, whereas we used the pseudo-R² of Nagelkerke for SAR. This allows us to tease out the effects of each factor and their interaction by building the following models :

1) M1 = model with all variables
2) M2 = model with all variables minus the variable of interest

The R² attributed solely to the variable of interest is the R² of M1 minus M2.

Then, we can tease out the spatial component from the effects of environmental predictors by building the following models :

1) M3 = model with all variables (with the spatial component)
2) M4 = model with all variables (without the spatial component)

In the case of SAR, the spatial component cannot be removed directly, so M4 was a linear model. The R² of Nagelkerke is equivalent to the classical R² of a linear regression (Nagelkerke, 1991), so we used this latter for M4. To test for the significance of spatial structure, we performed a Moran test on residuals of M4.

In the case of GDM, the spatial structure is directly included as a predictor by a matrix of geographic distances between sites/cells. Significance can be assessed by permutation tests as for other predictors.

Due to approximations in a few cases, the total R² of a complete model may appear to be slightly below the total of environmental and spatial parts of variance explained. In the opposite case (R² greater than the sum of variances explained by environment and spatial structure), it implies that there are some interaction parts among components, i.e. variance explained jointly by several factors (these parts are not shown here). Also, some fractions of variance can be negative in case of non-linear dependencies and are set to zero for the sake of interpretation.

**Appendix D.** Details about the sampling at each spatial resolution and test of the robustness of our analyses to sampling biases related to the spatial resolution.

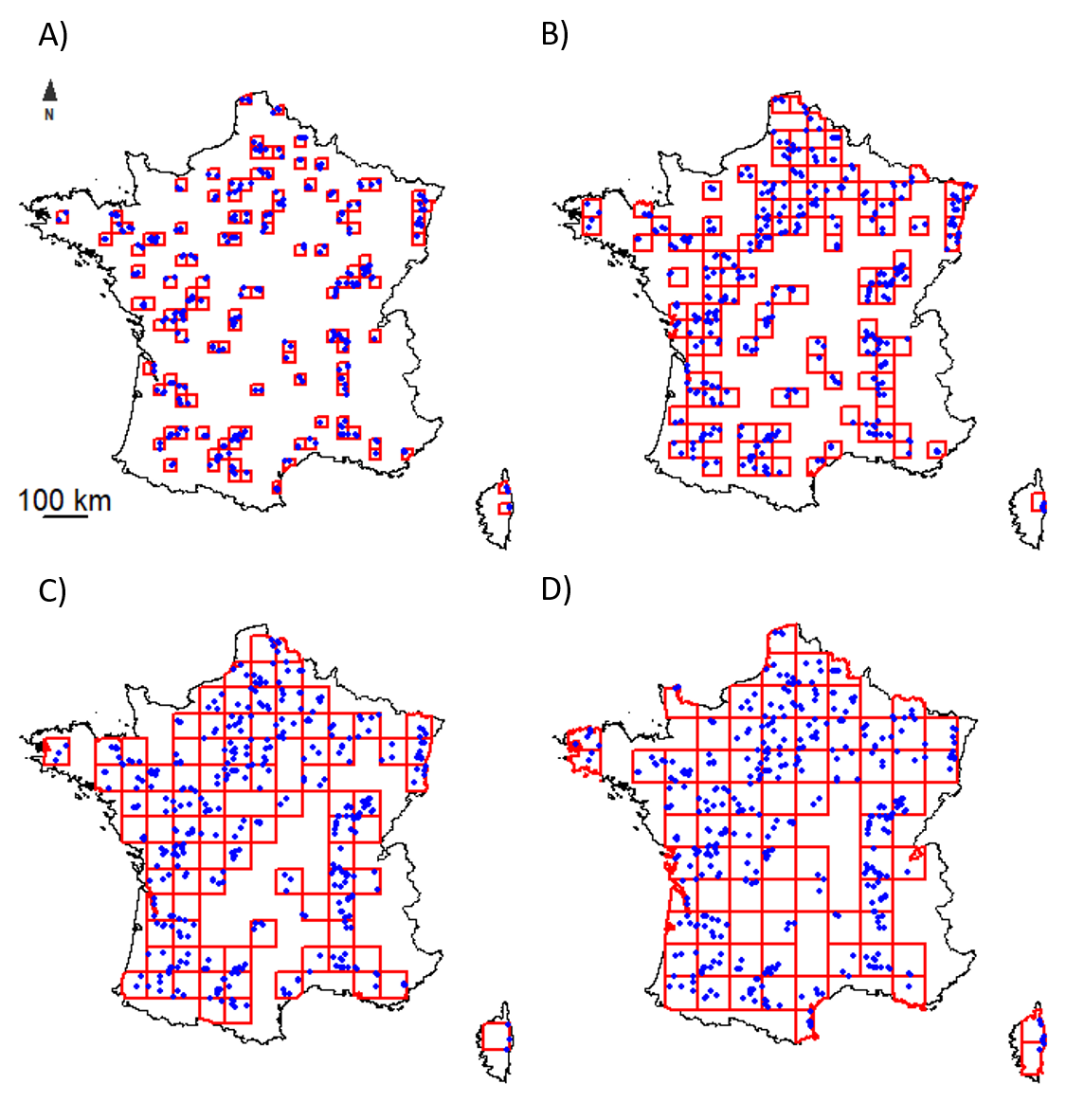

**Fig. D.1.** Distribution maps of the field margins (blue points) surveyed in grids (red) of different resolutions. A) 25 km, B) 40 km, C) 60 km and D) 75 km. The grid position has been shifted three times in analysis for each resolution. We shifted the initial grid position towards the northeast by half the length of a cell and towards the northwest by a quarter the length of a cell.

To help the reader understand our selection of resolutions, we clarify the factors that influenced our decision. First, we set the minimum and maximum resolutions at the site-level and 75 km, respectively, as larger grid cells would have resulted in too few cells for analysis. A Mantel correlogram on composition and a variogram on species richness (at the site-resolution) revealed that spatial autocorrelation was significant up to about 40-50 km, so we found it useful to include some resolutions below and above this threshold, while trying to keep resolution intervals more or less equally spaced. We wanted to include at least a couple of intermediate resolutions so that we made sure that the results were a logic progression between both ends, and not just some random variations due to a sampling / aggregation artifact. This is why we ended up with the 5 following resolutions: plot-level, 25, 40, 60 and 75 km. For simplicity, and because the 25 and 60 km resolutions only displayed minor differences with resolutions that preceded and followed them, we decided to present only the site-level, 40 km, and 75 km resolutions in the text.

Different biases can result from this analysis method built around the influence of spatial resolution. Indeed, the number of sites differs between cells and are not uniformly distributed in the cell. This could potentially lead to a lack of precision in cells containing fewer sites or sites that are not evenly distributed spatially. To test for this bias, we randomly drew 2 sites in each cell of the 25 km resolution, before running analyses and repeated the procedure 1000 times. We focused on the 25 km resolution as it is the most concerned by the low number of sites by cell (**Table D.1**). For each type of model (GDM and SAR), we plotted the distribution of parameters over these 1000 runs and compared it to the observed values. We can see that the bootstrap procedure induces little variation of the model parameters and that it is particularly true for SAR models (**Fig. D.2**).

**Table D.1.** Sample size for each resolution. Grid positions in the aggregated resolutions were shifted to obtain three different positions of the grid system. Means and standard deviations below reflect variations across these grids within each resolution.

| Spatial resolution | Site-level | 25 km | 40 km | 60 km | 75 km |
| --- | --- | --- | --- | --- | --- |
| Mean number of sites by cell |  | 2.7 ± 0,1 | 3.3 ± 0,1 | 4.6 ± 0,1 | 5.8 ± 0,1 |
| Mean number of sites | 462 | 306 ± 10 | 387 ± 9 | 418 ± 12 | 428 ± 1 |
| Mean number of cells |  | 87 ± 7 | 105 ± 3 | 86 ± 2 | 71 ± 2 |

A) **Composition (GDM)**

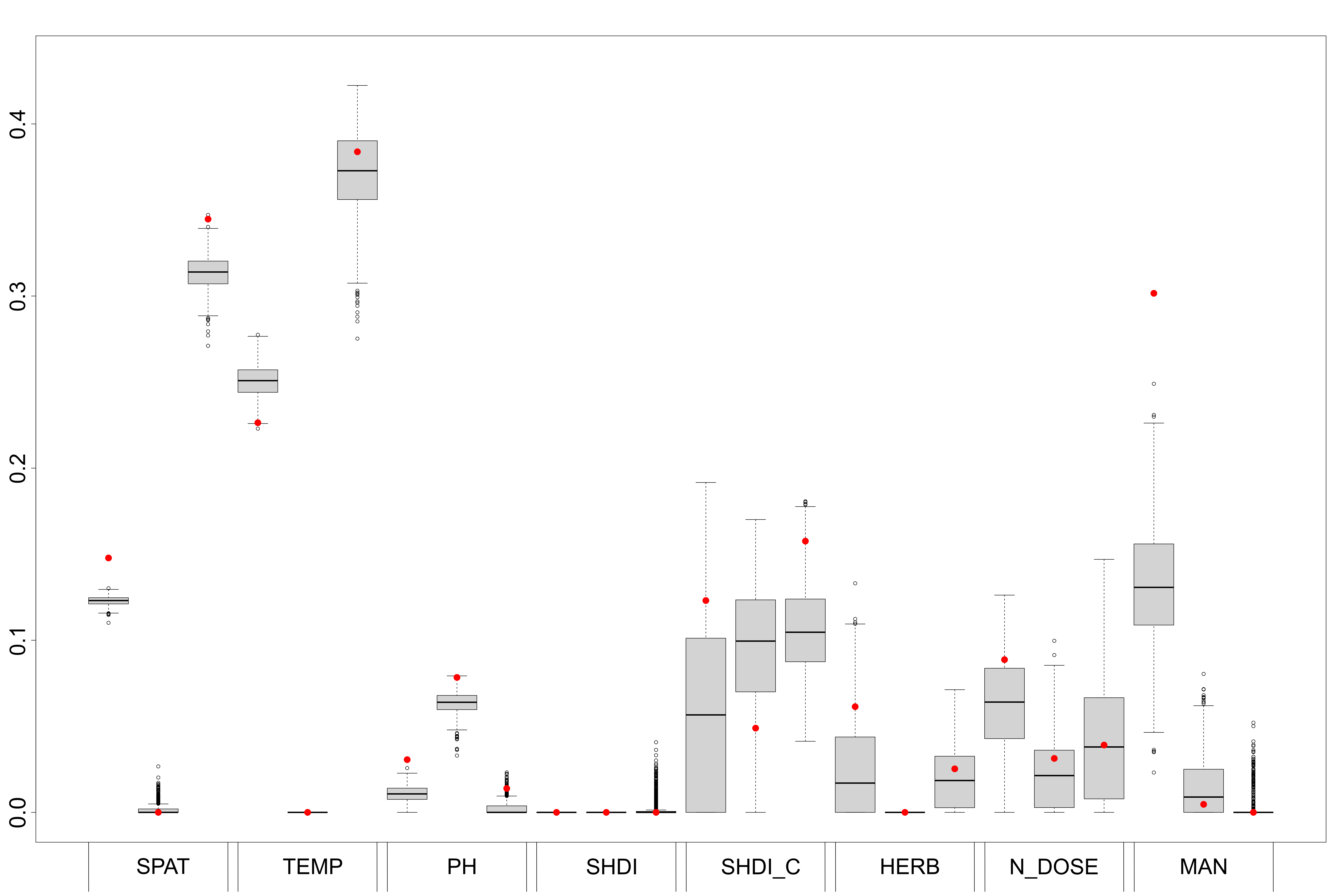

B) **Species richness (SAR)**

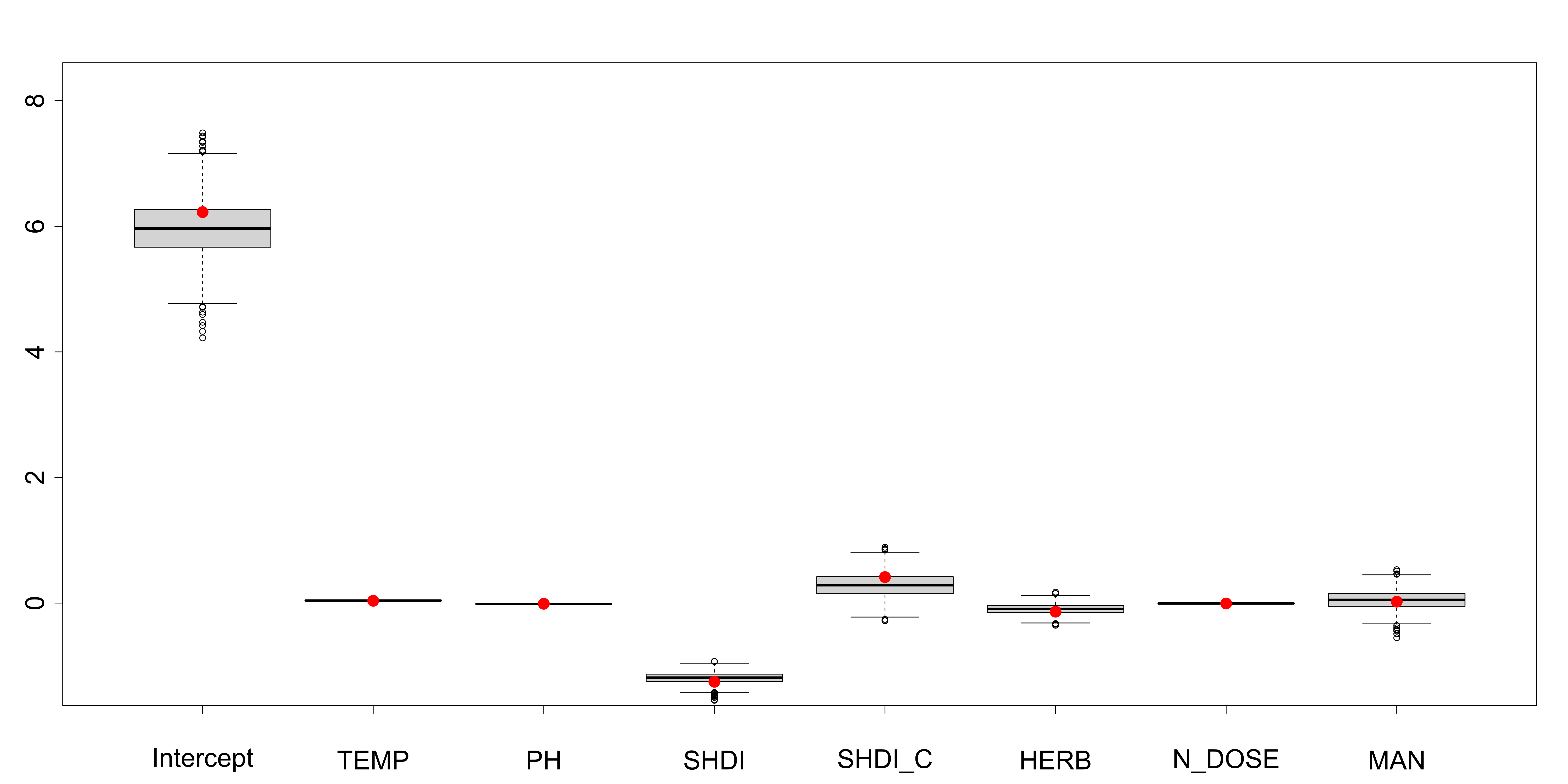

**Fig. D.2.** Variation of the A) GDM and B) SAR parameters over the 1000 resamplings. The red points are the observed coefficients. See **Table 1** for coefficient abbreviations. A) For GDM, each explanatory factor has three parameters, corresponding to three I-spline basis functions (see Ferrier et al., 2007).

A second bias that we wanted to test is the influence of the number of cells on the coefficients and R²/ explained deviance of the models. Indeed, the R² of a model can be artefactually inflated when the sampling power is too low, which is a problem when we aim to compare spatial resolutions with different number of spatial units. Consequently, we randomly draw 71 cells at each resolution (except for the coarsest one) as it is the mean number of cells for the resolution with the least one (75 km resolution). We performed analyses for the resolutions 0, 25, 40 and 60 km and repeated the procedure 1000 times. For each type of model (GDM and SAR), we plotted the distribution of the global R² / explained deviance over these 1000 runs and compared it to the observed values. We can see that the bootstrap procedure induces little variation of the total explained variance with a slight bias towards overestimation (**Fig. D.3**). However, the relative differences in explained variance between spatial resolutions are preserved with these resamplings, which comforts us in the fact that we can reliably compare spatial resolutions despite a different number of samples.

A) **Composition (GDM)**

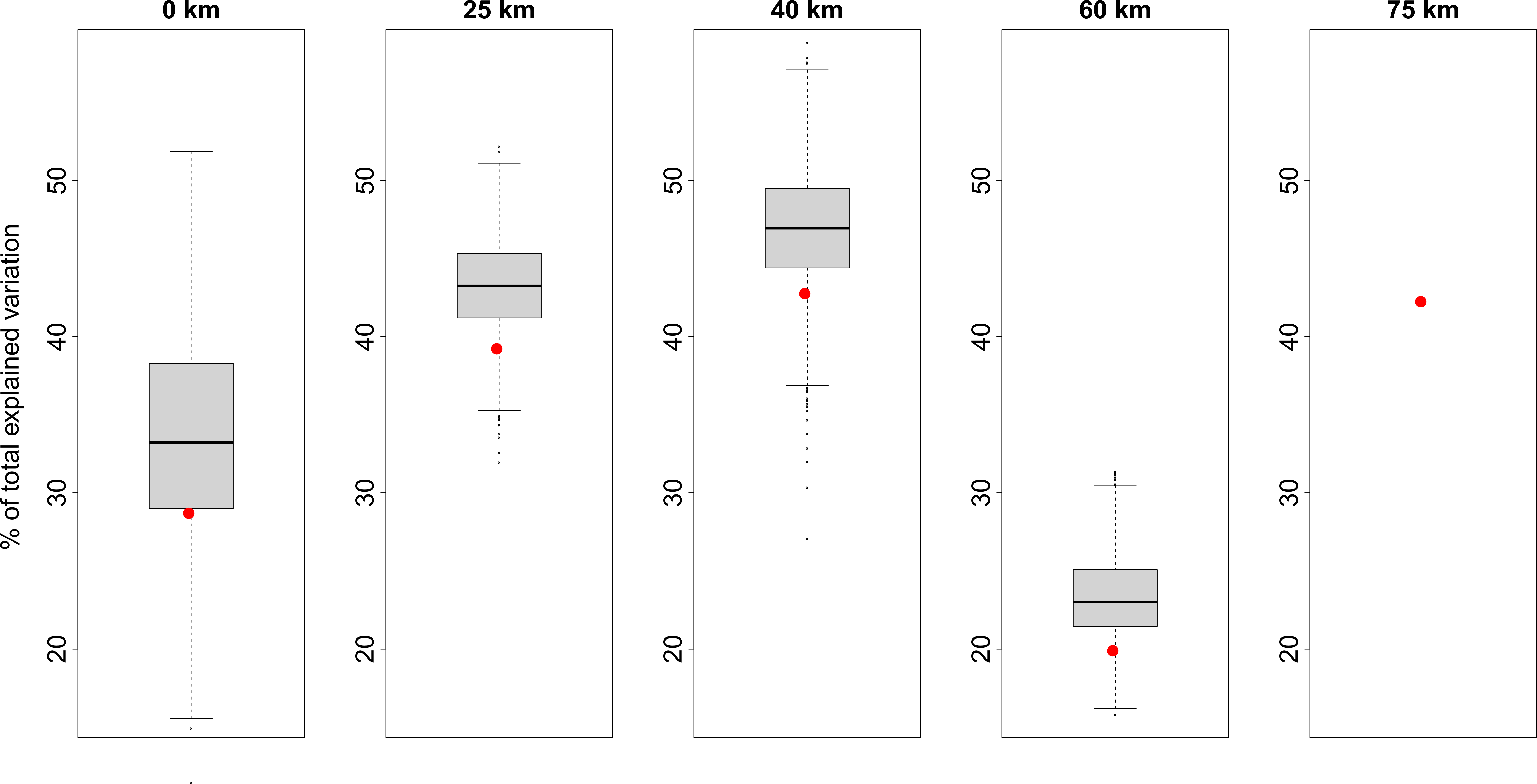

B) **Species richness (SAR)**

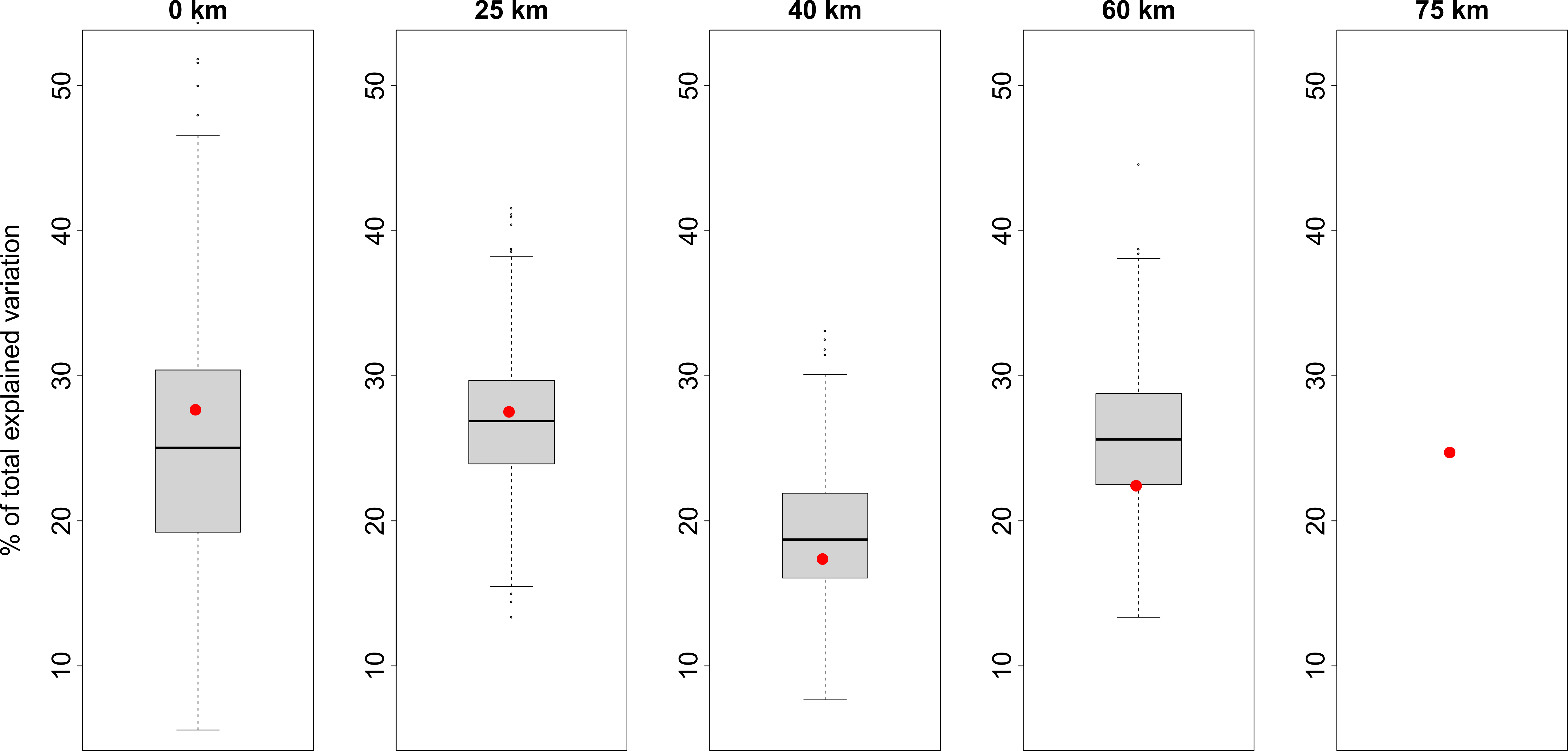

**Fig. D.3.** Variation of the A) GDM and B) SAR explained variance over the 1000 resamplings. The red points are the observed R² / explained deviance.

Another potential shortcoming was that the means computed within aggregated grid cells might not fully represent the regional pool or agricultural region. Indeed, the monitoring sites were stratified to represent the main crops and practices at the national level and not at the landscape or regional level. Checking the representativeness of the sites with the regional context is thus key to assess the relevance of the results. Unfortunately, we were unable to perform this validation with the available dataset.

**Appendix E**. Detailed outputs of GDM and SAR.

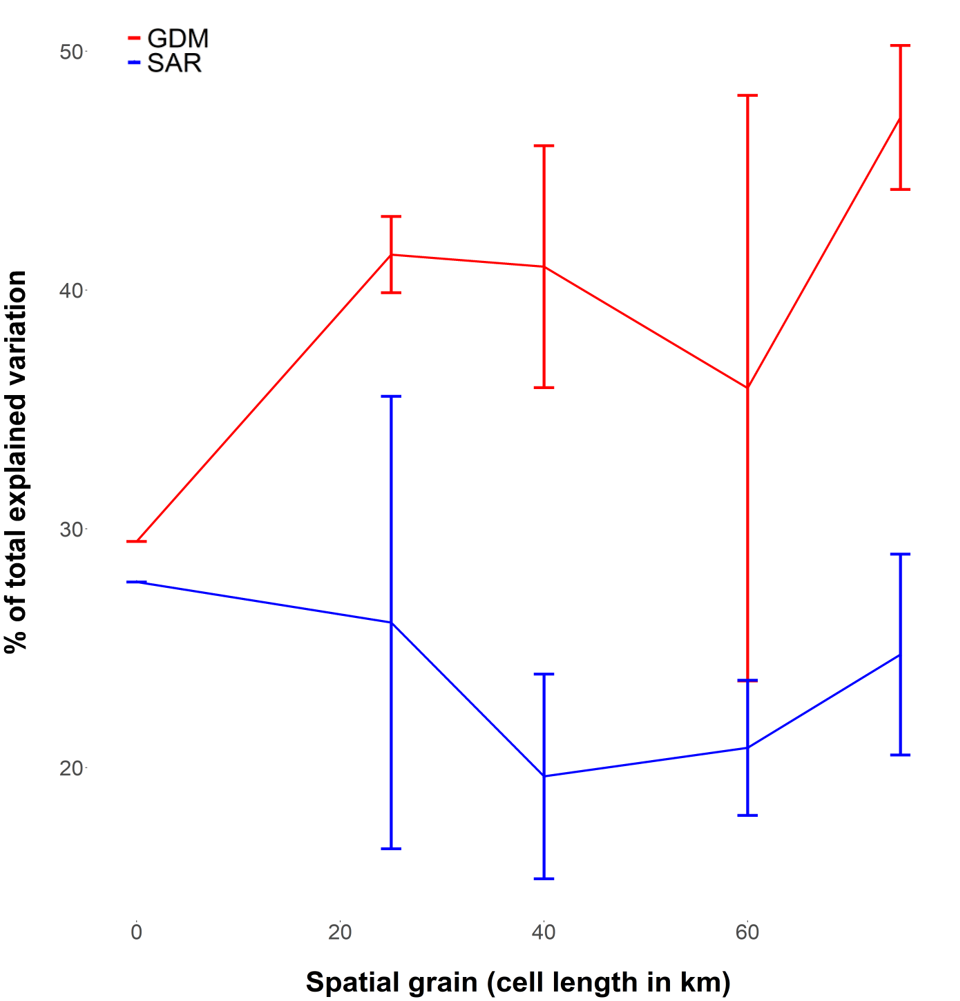

**Fig. E.1.** Total percentage of explained variance according to the spatial resolution of analyses (for GDM and SAR). Standard deviations were computed from the three resamplings of grids.

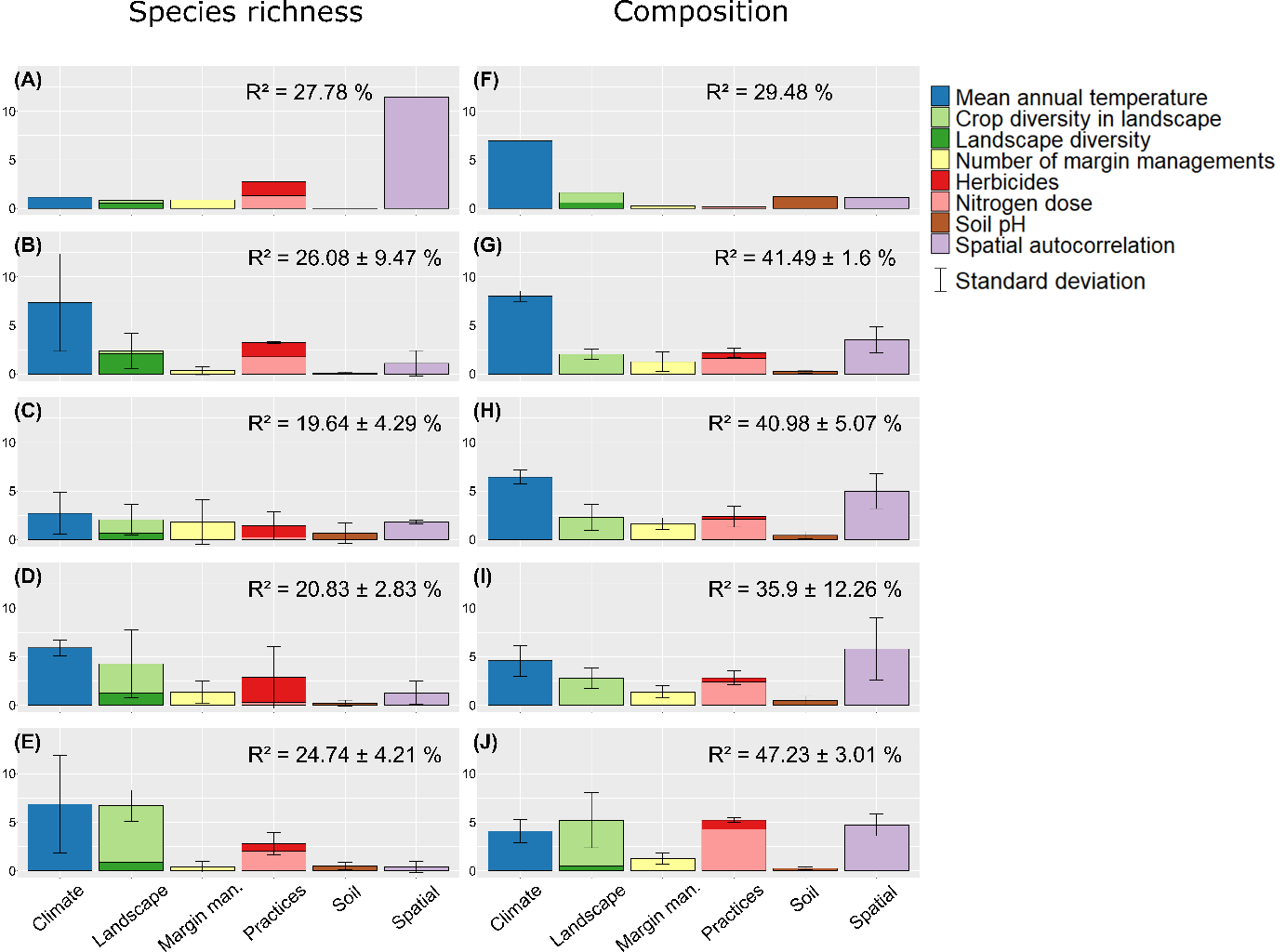

**Fig. E.2.** Percentage of explained variance of species richness and composition by each factor for each spatial resolution. (A, F) 0 km; (B, G) 25 km; (C, H) 40 km; (D, I) 60 km; (E,J) 75 km. Standard deviations are computed from the three resamplings of grids.

**Table E.1.** Significance of each predictor (in row) for each spatial resolution (in column, in km). For species richness in A), the direction of the relationship is indicated by +/-. The number of +/-/* indicates the number of times the predictor was significant over the three grid shifts. We only report significant relationships with p-values in brackets (the average p-value when significant in multiple grids).

**A) Species richness**

|  | 0 | 25 | 40 | 60 | 75 |
| --- | --- | --- | --- | --- | --- |
| Temperature | + (0.004) | ++ (0.002) | + (0.010) | +++ (0.012) | ++ (0.011) |
| Ph |  |  |  |  |  |
| Landscape diversity |  | - (0.041) |  |  |  |
| Crop diversity |  |  |  | ++ (0.027) | +++ (0.017) |
| Herbicides | - (0.002) | - (0.048) | - (0.031) | - (0.006) |  |
| Nitrogen dose | - (0.004) |  |  |  | - (0.044) |
| Margin management | + (0.018) |  | + (0.019) |  |  |
| Spatial structure | (< 0.001) |  |  |  |  |

**B) Composition**

|  | 0 | 25 | 40 | 60 | 75 |
| --- | --- | --- | --- | --- | --- |
| Temperature | (< 0.001) | *** (< 0.001) | *** (< 0.001) | *** (0.013) | *** (0.020) |
| Ph | (< 0.001) |  |  |  |  |
| Landscape diversity | (< 0.001) |  |  |  |  |
| Crop diversity | (< 0.001) | ** (< 0.001) | ** (< 0.001) | ** (0.010) | *** (0.020) |
| Herbicides |  |  |  |  |  |
| Nitrogen dose |  | *** (0.030) | ** (< 0.001) | ** (0.020) | *** (0.010) |
| Margin management |  | * (0.020) | * (< 0.001) |  |  |
| Spatial structure | (< 0.001) | *** (< 0.001) | *** (< 0.001) | *** (< 0.001) | *** (< 0.001) |

**Table E.2.** Mean percentages ± standard deviations of explained variance of A) species richness and B) composition by each factor (in row) for each spatial resolution (in column, in km). Means and standard deviations are computed from the three grids. We included the portion of variance that results from the combined influence of multiple factors, which is commonly referred to as "Interactions."

**A) Species richness**

|  | 0 | 25 | 40 | 60 | 75 |
| --- | --- | --- | --- | --- | --- |
| Temperature | 1.15 | 7.35 ± 5.02 | 2.71 ± 2.15 | 5.90 ± 0.82 | 6.85 ± 5.01 |
| Ph | 0 | 0.09 ± 0.12 | 0.66 ± 1.05 | 0.21 ± 0.3 | 0.51 ± 0.38 |
| Landscape diversity | 0.53 | 2.08 ± 1.42 | 0.67 ± 0.65 | 1.21 ± 0.98 | 0.86 ± 0.95 |
| Crop diversity | 0.31 | 0.30 ± 0.44 | 1.37 ± 0.97 | 3.05 ± 2.65 | 5.83 ± 1.14 |
| Herbicides | 1.47 | 1.47 ± 1.46 | 1.21 ± 1.53 | 2.56 ± 3.29 | 0.80 ± 0.81 |
| Nitrogen dose | 1.28 | 1.76 ± 1.53 | 0.23 ± 0.16 | 0.29 ± 0.3 | 1.99 ± 1.95 |
| Margin management | 0.87 | 0.33 ± 0.41 | 1.83 ± 2.3 | 1.32 ± 1.16 | 0.42 ± 0.55 |
| Spatial structure | 11.43 | 1.09 ± 1.32 | 1.8 ± 0.19 | 1.28 ± 1.22 | 0.38 ± 0.59 |
| Interactions | 10.74 | 11.61 ± 3.05 | 9.16 ± 6.49 | 5.02 ± 4.49 | 7.11 ± 6.93 |
| R² | 27.78 | 26.08 ± 9.47 | 19.64 ± 4.29 | 20.83 ± 2.83 | 24.74 ± 4.21 |

**B) Composition**

|  | 0 | 25 | 40 | 60 | 75 |
| --- | --- | --- | --- | --- | --- |
| Temperature | 6.93 | 8.01 ± 0.53 | 6.44 ± 0.7 | 4.56 ± 1.59 | 4.08 ± 1.22 |
| Ph | 1.23 | 0.25 ± 0.15 | 0.43 ± 0.38 | 0.46 ± 0.45 | 0.25 ± 0.15 |
| Landscape diversity | 0.59 | 0 ± 0 | 0.01 ± 0.02 | 0.04 ± 0.04 | 0.51 ± 0.59 |
| Crop diversity | 1.05 | 2.04 ± 0.52 | 2.29 ± 1.37 | 2.73 ± 1.06 | 4.69 ± 2.27 |
| Herbicides | 0.04 | 0.55 ± 0.56 | 0.29 ± 0.26 | 0.40 ± 0.29 | 0.95 ± 0.54 |
| Nitrogen dose | 0.14 | 1.63 ± 0.30 | 2.07 ± 0.91 | 2.42 ± 0.44 | 4.28 ± 0.79 |
| Margin management | 0.22 | 1.26 ± 1.01 | 1.65 ± 0.59 | 1.37 ± 0.64 | 1.28 ± 0.57 |
| Spatial structure | 1.09 | 3.55 ± 1.34 | 4.98 ± 1.79 | 5.79 ± 3.22 | 4.73 ± 1.13 |
| Interactions | 18.2 | 24.2 ± 3.31 | 22.81 ± 4.47 | 18.13 ± 6.26 | 26.48 ± 6.92 |
| Explained deviance | 29.48 | 41.49 ± 1.6 | 40.98 ± 5.07 | 35.9 ± 12.26 | 47.23 ± 3.01 |

**Table. E.3.** Percentages of explained variance of A) species richness and B) composition by each factor (in row) for each biogeographic region and for national extent (in column, see **Fig. 1** for abbreviations of region names). We included the portion of variance that results from the combined influence of multiple factors, which is commonly referred to as "Interactions."

**A) Species richness**

|  | BPN | BPS | MA | ZM | ZNE | ZSO | France |
| --- | --- | --- | --- | --- | --- | --- | --- |
| Temperature | 0.70 | 1.13 | 0.18 | 0 | 2.74 | 6.25 | 1.15 |
| Ph | 0.86 | 2.44 | 0 | 0 | 2.01 | 3.41 | 0 |
| Landscape diversity | 6.49 | 1.23 | 0 | 2.95 | 3.13 | 0.49 | 0.53 |
| Crop diversity | 0.96 | 1.55 | 1.53 | 1.03 | 2.14 | 1.35 | 0.31 |
| Herbicides | 5.95 | 1.27 | 0 | 0 | 3.54 | 1.20 | 1.47 |
| Nitrogen dose | 0.64 | 2.79 | 0.02 | 12.88 | 3.02 | 1.65 | 1.28 |
| Margin management | 6.58 | 1.25 | 0 | 7.07 | 2.07 | 4.46 | 0.87 |
| Spatial structure | 0.64 | 2.58 | 1.65 | 0.02 | 10.36 | 11.4 | 11.43 |
| Interactions | 1.00 | 8.32 | 11.67 | 14.84 | 0 | 2.86 | 10.74 |
| R² | 23.82 | 22.56 | 15.05 | 38.79 | 26.81 | 33.07 | 27.78 |

**B) Composition**

|  | BPN | BPS | MA | ZM | ZNE | ZSO | France |
| --- | --- | --- | --- | --- | --- | --- | --- |
| Temperature | 0.79 | 1.36 | 0.13 | 6.19 | 0.88 | 0.74 | 6.93 |
| Ph | 0.26 | 1.90 | 0.21 | 0.84 | 0.08 | 0.41 | 1.23 |
| Landscape diversity | 1.86 | 0.02 | 2.17 | 2.23 | 0.75 | 0.07 | 0.59 |
| Crop diversity | 1.05 | 0 | 0.79 | 1.65 | 1.04 | 1.04 | 1.05 |
| Herbicides | 1.12 | 0.08 | 0.81 | 0.01 | 1.54 | 0 | 0.04 |
| Nitrogen dose | 0 | 1.37 | 0.11 | 1.11 | 0.72 | 0 | 0.14 |
| Margin management | 0.62 | 0.17 | 6.87 | 5.58 | 0 | 0.98 | 0.22 |
| Spatial structure | 7.83 | 4.83 | 2.32 | 7.30 | 3.30 | 9.41 | 1.09 |
| Interactions | 9.53 | 3.51 | 4.64 | 19.67 | 2.29 | 4.00 | 18.19 |
| Explained deviance | 23.06 | 13.24 | 18.05 | 44.58 | 10.60 | 16.65 | 29.48 |

**Appendix F.** Range of values of explanatory variables within spatial resolutions and biogeographic regions.

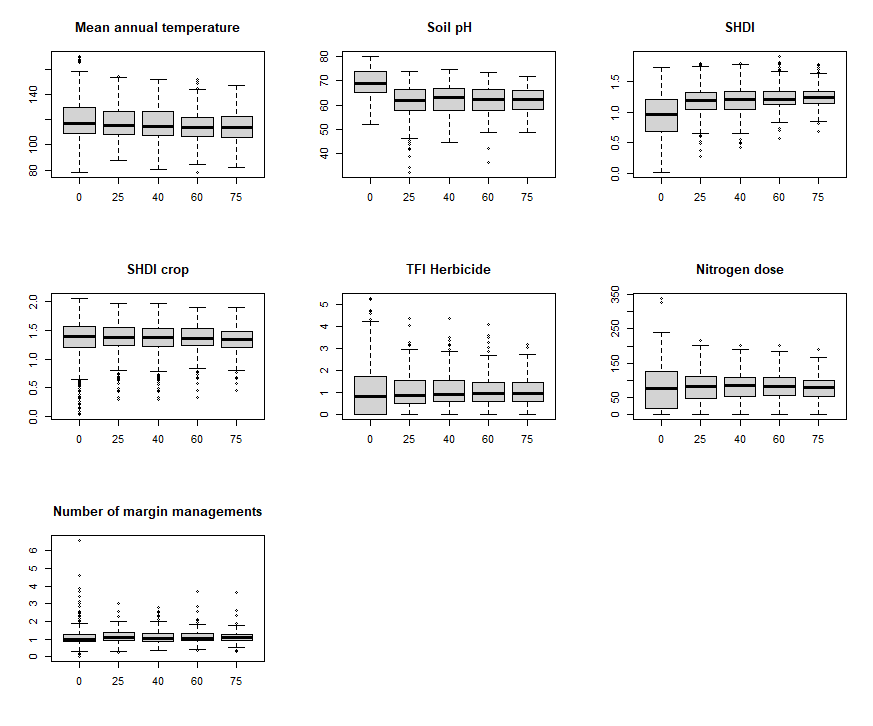

**Fig. F.1.** Boxplot of environmental predictors by spatial resolution (in km). See **Table 1** for units.

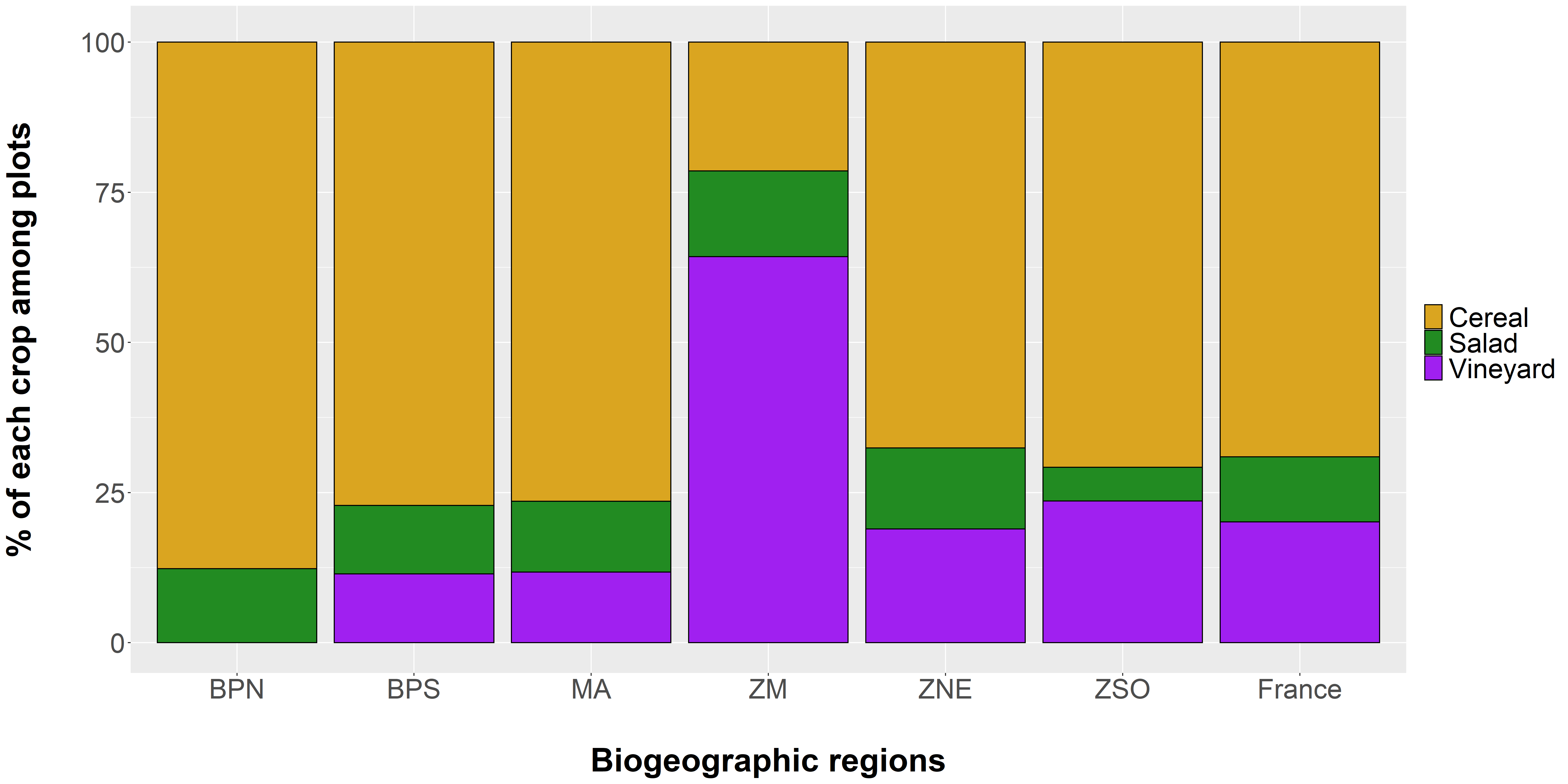

**Fig. F.2.** Percentages of each crop type among sites surveyed within each biogeographic region. See **Fig. 1** for abbreviations of region names.

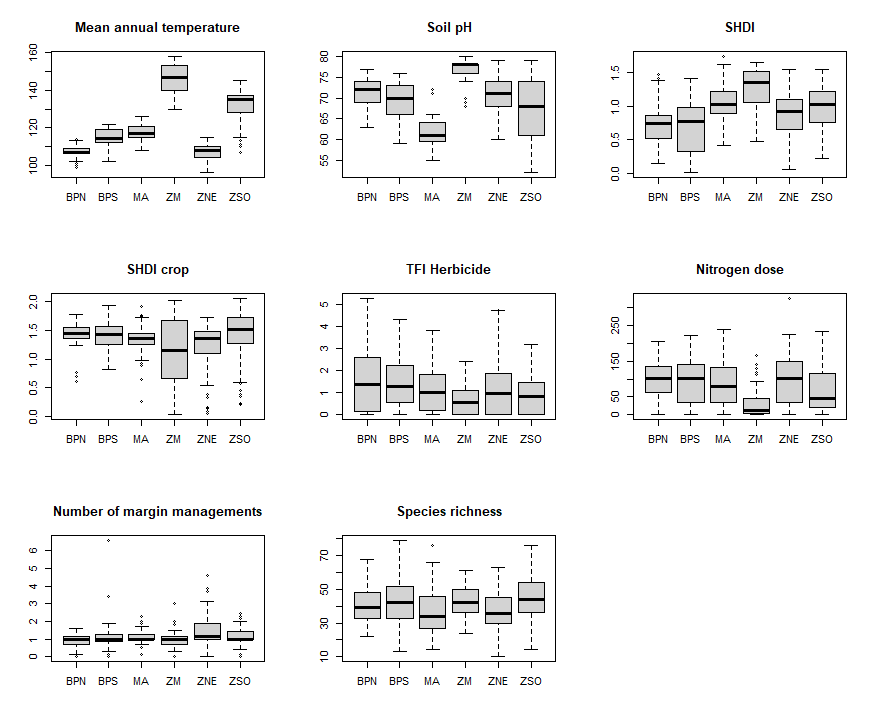

**Fig. F.3.** Boxplot of environmental predictors by biogeographic region. See **Fig. 1** for abbreviations of region names and **Table 1** for units.
